# Hepatocyte Rap1a Contributes To Obesity- and Statin-Associated Hyperglycemia

**DOI:** 10.1101/2022.06.25.497009

**Authors:** Yating Wang, Stefano Spolitu, John A. Zadroga, Amesh K. Sarecha, Lale Ozcan

## Abstract

Excessive hepatic glucose production contributes to the development of hyperglycemia and is a key feature of type 2 diabetes. Here, we report that activation of hepatic Rap1a suppresses gluconeogenic gene expression and glucose production, whereas Rap1a silencing stimulates them. Rap1a activation is suppressed in obese mouse liver and restoring its activity lowers blood glucose and improves glucose intolerance. As Rap1a’s membrane localization and activation depends on its geranylgeranylation, which is inhibited by statins, we found lower active-Rap1a levels in statin-treated hepatocytes and the human liver. Similar to Rap1a inhibition, statins stimulated hepatic gluconeogenesis and increased fasting blood glucose in obese mice. Geranylgeraniol treatment, which acts as the precursor for geranylgeranyl isoprenoids, restored Rap1a activity and improved statin-mediated glucose intolerance. Mechanistically, we show that Rap1a activation induces actin polymerization, which suppresses gluconeogenesis by Akt-mediated FoxO1 inhibition. Thus, Rap1a regulates hepatic glucose homeostasis, and blocking its activity, via lowering geranylgeranyl isoprenoids, contributes to statin-induced glucose intolerance.

## INTRODUCTION

A major cause of hyperglycemia in type 2 diabetes (T2D) is enhanced hepatic glucose production (HGP), which results from an increase in gluconeogenesis (Lin and Accili, 2011; Magnusson et al., 1992). Several factors, including the availability of substrates and an imbalance between glucagon and insulin action contribute to this process (Petersen et al., 2017). Glucagon plays a major role in hepatic gluconeogenesis, in part by increasing cyclic AMP (cAMP) and intracellular calcium levels, which result in increased expression of two key gluconeogenic enzymes, phosphoenol pyruvate carboxykinase (PCK1) and glucose 6 phosphatase (G6PC) (Ramnanan et al., 2011). A number of transcription factors and coactivators including, forkhead box O1 (FoxO1), cAMP response element-binding protein (CREB) and CREB-regulated transcription co-activator 2 (CRTC2) activate and stimulate the transcription of these gluconeogenic genes in T2D; however, little is known about the endogenous regulatory mechanisms that prevent aberrant activation of this pathway.

One of the major complications of T2D is to increase the risk for developing cardiovascular disease, and the majority of T2D patients use cholesterol-lowering statin drugs (Benjamin et al., 2018). By inhibiting Hmg-CoA reductase, the rate-limiting enzyme of the mevalonate pathway, statins decrease hepatic cholesterol synthesis and increase low-density lipoprotein (LDL) receptor (LDLR) levels, which results in enhanced LDL clearance from the plasma (Baigent et al., 2005; Goldstein and Brown, 2015). Although they are generally considered to be safe and well-tolerated, recent evidence from meta-analyses of major statin trials showed that statin therapy is associated with a 9-10% increased risk for new-onset T2D (Casula et al., 2017; Ridker et al., 2008; Sattar et al., 2010). The risk is dose-dependent and increased by close to 46% in older patients and patients with pre-existing risk factors for T2D (Crandall et al., 2017; Preiss et al., 2011; Ridker et al., 2012). These results prompted US Food and Drug Administration (FDA) to approve labeling changes in statins to include a warning about the possibility of increased blood sugar (FDA, 2012). Interestingly, loss-of-function mutations in *HMGCR*, which encodes Hmg-CoA reductase, are also associated with increased incidence of new-onset T2D, suggesting that the diabetogenic effect of statins is “on-target” (Ference et al., 2016; Swerdlow et al., 2015). Despite these findings, the cellular and molecular mechanisms that link statins to increased T2D risk are largely unknown. Besides lowering plasma LDL cholesterol (LDL-C) levels, statins have additional actions, which involve inhibition of the mevalonate pathway. Mevalonate is the precursor for the generation of farnesyl pyrophosphate (FPP) and geranylgeranyl pyrophosphate (GGPP) isoprenoids that are required for prenylation and activation of a subset of proteins, including the small GTPase, Ras-related protein 1A (Rap1a).

Rap1a belongs to the Ras superfamily of GTPases and cycles between an inactive GDP-bound form and an active GTP-bound form (Gloerich and Bos, 2011; Kitayama et al., 1989; Wittchen et al., 2011). Post-translational covalent attachment of geranylgeranyl isoprenoid chain to Rap1a’s C-terminal cysteine residue results in its membrane localization, where exchange protein directly activated by cAMP 2 (Epac2) activates Rap1a via GTP loading (Bos et al., 2007; de Rooij et al., 1998). GTPase-activating proteins, such as Rap1GAP, stimulate GTP hydrolysis and thereby inactivate Rap1a. We previously reported a role for Rap1a in metabolic disease and showed that activation of Rap1a lowers plasma LDL-C via decreasing proprotein convertase subtilisin-kexin type 9 (PCSK9) (Spolitu et al., 2019). However, much less is known about the contribution of hepatic Epac2 or Rap1a to glucose homeostasis.

In the present study, we explored the role of Rap1a in liver glucose metabolism and show that hepatic Rap1a suppresses gluconeogenesis and glucose production via regulating actin polymerization and Akt-mediated FoxO1 activation. Hepatic Rap1a activity is significantly lower in obese mice, and overexpression of a constitutively active mutant form of Rap1a in the liver lowers blood glucose and improves glucose intolerance. Given that geranylgeranylation is required for Rap1a to become fully activated, we show that Rap1a activity is also decreased in statin-treated primary hepatocytes and human liver samples from statin users. Accordingly, statin treatment stimulates glucose production *in vitro* and further increases fasting blood glucose and worsens glucose intolerance in obese mice. Adding back geranylgeranyl isoprenoids restores hepatic Rap1a activity and improves hyperglycemia. These findings identify Rap1a as an important mediator of obesity- and statin-induced glucose intolerance.

## RESULTS

### Inhibition of Epac2 or Rap1a Stimulates Gluconeogenesis and Increases HGP

A recent GWAS study demonstrated that variants in *RAPGEF4*, which encodes EPAC2, are associated with fasting blood glucose in East Asians; however, the functional significance of this finding is not known (Hwang et al., 2015). To test whether Epac2 and its downstream effector, Rap1a, contribute to hepatic glucose metabolism, we tested their effects on the expression of gluconeogenic genes, *G6Pc* and *Pck1.* Upon stimulation of freshly isolated primary mouse hepatocytes with forskolin, a glucagon mimetic and potent adenylate cyclase activator, and dexamethasone, which downregulates cAMP-phosphodiesterase (Manganiello and Vaughan, 1972), we observed that expression of *G6Pc* and *Pck1* were further increased in WT hepatocytes treated with siRNAs against Epac2 or Rap1a (**Figures 1A, 1C, and 1D**). Accordingly, glucagon treatment resulted in an increase in *G6pc* and *Pck1* mRNA levels in Rap1a-deficient hepatocytes versus control (**Figure 1F**). Similar data were obtained in a non-cancerous mouse hepatocyte cell line, AML-12 cells (**Figures S1A and S1B**). Consistent with an increase in the expression of gluconeogenic genes, we observed a further increase in glucose release in cells lacking Epac2 or Rap1a after forskolin and dexamethasone stimulation (**Figures 1B and 1E**). To assess the functional role of Rap1a in hepatic glucose metabolism *in vivo*, we deleted liver Rap1a by injecting diet-induced obese (DIO) *Rap1a*^fl/fl^ mice with adeno-associated virus-8 encoding Cre recombinase driven by the hepatocyte-specific thyroxine-binding globulin promoter (AAV8-TBG-Cre), and AAV8-TBG-Gfp-treated *Rap1a*^fl/fl^ mice served as controls. The TBG-Cre treatment successfully silenced Rap1a in the liver (**Figure 1G**) without altering body weight (**Figure 1H**). In line with the *in vitro* data, DIO mice that lack hepatocyte

**Figure 1.**
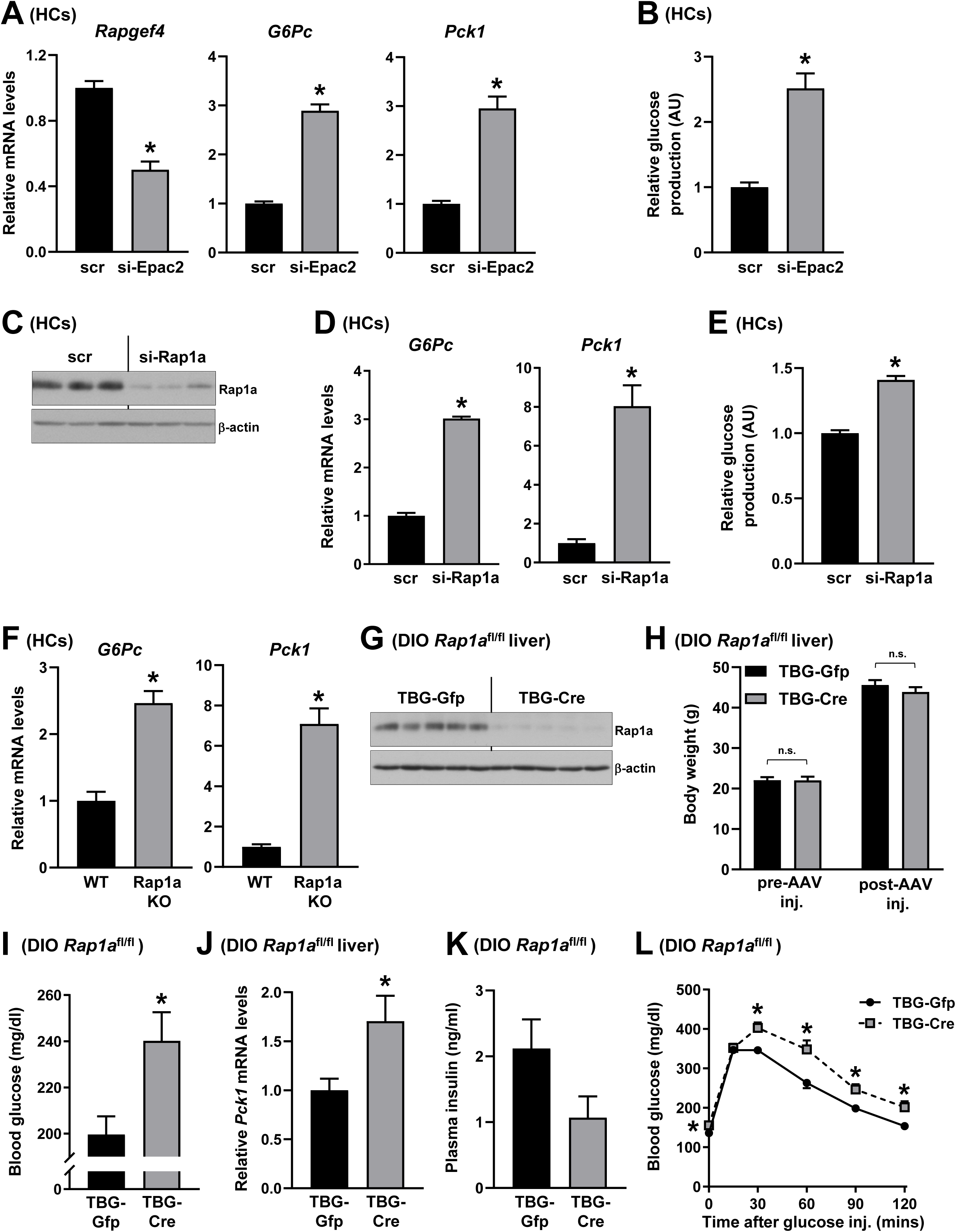
Inhibition of Epac2 or Rap1a Stimulates Gluconeogenesis and Increases Hepatic Glucose Production. **(A)** *Rapgef4* (Epac2), *G6Pc* and *Pck1* mRNA levels were analyzed from forskolin and dexamethasone-treated primary mouse hepatocytes (HCs) that were transfected with scrambled RNA (scr) or siRNA against Epac2 (si-Epac2) (n= 4 wells of cells/group, mean ± SEM, *p < 0.05). **(B)** Same as in (A), except that glucose production was measured (n= 4 wells of cells/group, mean ± SEM, *p < 0.05). **(C-E)** Rap1a and β-actin levels (A), *G6Pc* and *Pck1* mRNA (B), and glucose production (C) from forskolin and dexamethasone-treated primary mouse hepatocytes that were transfected with scrambled RNA (scr) or siRNA against Rap1a (si-Rap1a) (n= 3-4 wells of cells/group, mean ± SEM, *p < 0.05). **(F)** *G6Pc* and *Pck1* mRNA levels were measured from glucagon-treated WT or Rap1a KO primary mouse hepatocytes (n= 6 wells of cells/group, mean ± SEM, *p < 0.05). **(G-L)** Hepatic Rap1a and β-actin levels (G), body weight before and after AAV injection (H), 5 h fasting blood glucose (I), liver *Pck1* mRNA levels (J), 5 h fasting plasma insulin levels (K), and glucose tolerance test (L) from DIO *Rap1a*^fl/fl^ mice that were injected with adeno-associated viruses containing either hepatocyte-specific TBG-Cre recombinase (TBG-Cre) or the control vector (TBG-Gfp) (n= 5-8 mice/group, mean ± SEM, *p < 0.05, n.s., non-significant).

Rap1a had higher fasting blood glucose levels (**Figure 1I**) and hepatic *Pck1* expression (**Figure 1J**) without a change in plasma insulin levels (**Figure 1K**). Consistent with these results, hepatocyte Rap1a-deficient mice also showed greater blood glucose excursions than control mice upon glucose challenge (**Figure 1L**). We next evaluated the effect of activated hepatic Rap1a by treating primary hepatocytes with a plasmid encoding a constitutively active Rap1a mutant (CA-Rap1a). CA-Rap1a possesses an amino acid substitution, Q63E, which results in constitutive GTP-binding, and thus the mutated form of Rap1a is always active and unable to be downregulated by Rap1GAP (Spolitu *et al*., 2019; Wittchen *et al*., 2011). Compared with control plasmid-treated cells, CA-Rap1a-overexpressing cells had lower *G6Pc* and *Pck1* mRNA expression and showed 50% decrease in glucose production upon forskolin and dexamethasone treatment (**Figures S1C and S1D**). These results suggested that activation of Epac2 and Rap1a suppress gluconeogenesis.

To further evaluate the physiologic relevance of Rap1a mediated gluconeogenesis suppression, we checked hepatic Rap1a activity during the transition from a fed to fasting state in WT mice. Consistent with an increase in hepatic gluconeogenic gene expression, we found that GTP-bound, active-Rap1a levels in the liver were lower after 6 hours of fasting (**Figure S2A**), which remained low after 14 and 24 hours of fasting (**Figures S2B and S2C**). Fasting, however, did not increase the levels of hepatic phosphorylated CREB as reported before (Stern et al., 2019). These data support the hypothesis that Rap1a negatively regulates gluconeogenesis, and the major function of Rap1a in hepatic glucose metabolism is to inhibit aberrant activation of HGP.

### Activation of Rap1a is Suppressed in Obese Mice Liver, and Restoring Rap1a Activity Lowers Blood Glucose and Improves Glucose Intolerance in Obese Mice

Because Rap1a activation lowers gluconeogenesis and glucose production (Figure S1C and S1D), we next asked if Rap1a is inhibited in mouse models of diabetes and insulin resistance where gluconeogenesis and HGP are elevated. In genetically obese and diabetic *db/db* mice liver, we found that the GTP-bound, active Rap1a levels were significantly decreased as compared to heterozygous *db/+* controls (**Figure 2A**). Similar data were obtained in the livers of DIO mice versus their low-fat-fed controls (**Figure 2B**). We then restored hepatic Rap1a activity by treating *db/db* mice with AAV8-TBG-CA-Rap1a, which specifically expresses the constitutively active Rap1a in hepatocytes. Despite similar body weight (**Figure 2C**), we observed that overexpressing CA-Rap1a in *db/db* mice liver lowered fasting blood glucose (**Figure 2D**), improved glucose intolerance (**Figure 2E**) and lowered hepatic gluconeogenic gene expression (**Figure 2F**), without affecting plasma insulin levels (**Figure 2G**). Similar results were obtained when DIO mice were treated with adeno-CA-Rap1a (**Figures 2H-2L**). These combined data provide evidence that Rap1a regulates gluconeogenesis and HGP and contributes to glucose homeostasis *in vivo*.

**Figure 2.**
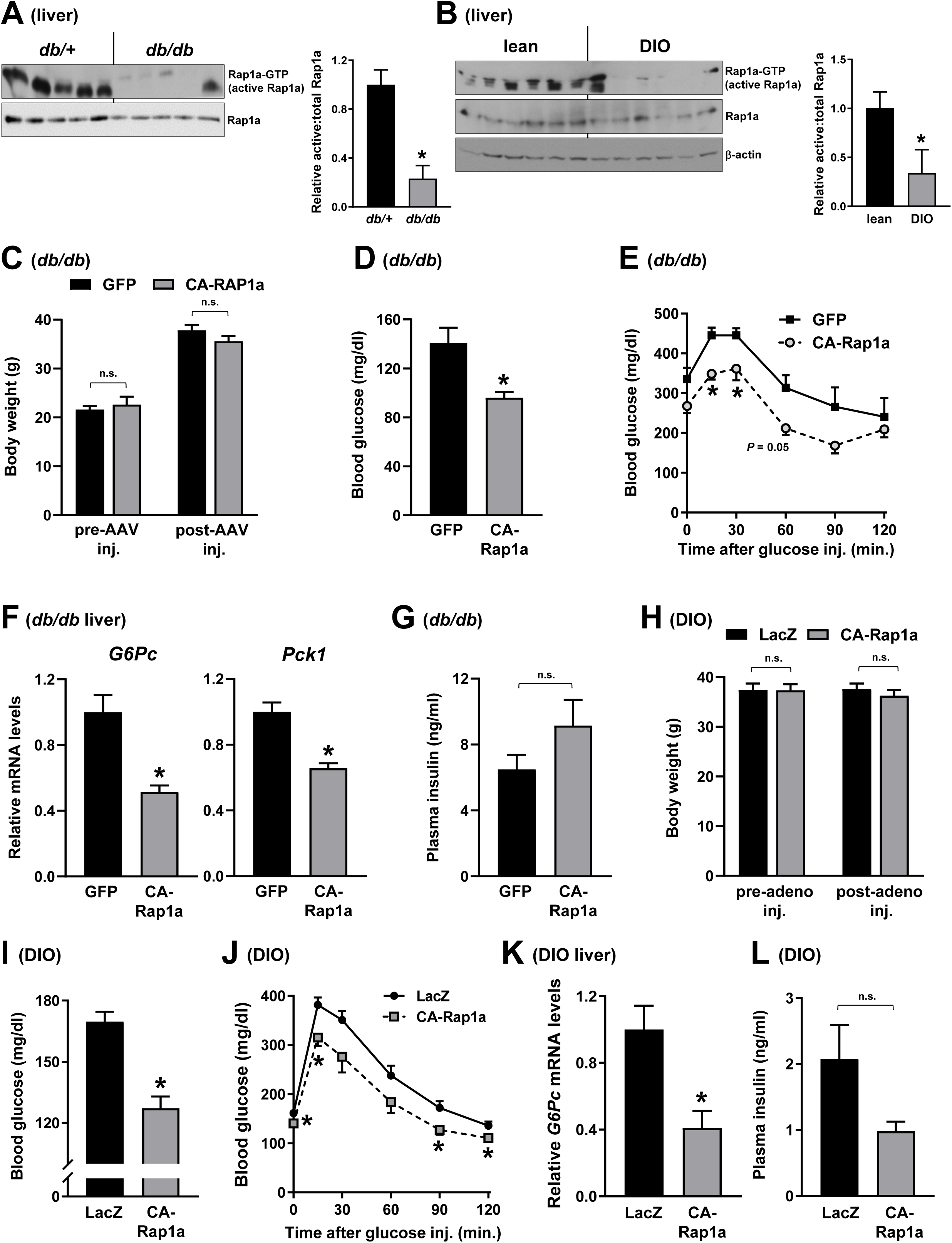
Activation of Rap1a Is Suppressed in Obese Mice Liver and Restoring Rap1a Activity Lowers Blood Glucose and Improves Glucose Intolerance in Obese Mice. **(A)** Livers from *db/db* and control (*db/+*) mice were assayed for GTP-bound (active) Rap1a and total Rap1a. Densitometric quantification of the immunoblot data is shown in the bar graph (n= 5 mice/group, mean ± SEM, *p < 0.05). **(B)** Livers from diet-induced obese (DIO) mice and their low-fat-fed controls (lean) were assayed for GTP-bound (active) Rap1a, total Rap1a and β-actin. Densitometric quantification of the immunoblot data is shown in the bar graph (n= 6 mice/group, mean ± SEM, *p < 0.05). **(C-G)** Body weight before and after AAV injection (C), overnight fasting blood glucose (D), glucose tolerance test (E), liver *G6Pc* and *Pck1* mRNA expression levels (F), and overnight fasting plasma insulin levels (G) from *db/db* mice that were injected with adeno-associated viruses (AAV) containing either hepatocyte-specific CA-Rap1a (constitutively active Rap1a) or the control vector (Gfp) (n= 4-7 mice/group, mean ± SEM, *p < 0.05, n.s., non-significant). **(H-L)** Body weight before and after adenovirus injection (H), fasting blood glucose (I), glucose tolerance test (J), and liver *G6Pc* mRNA expression levels (K), and 5 h fasting plasma insulin (L) from DIO mice that were injected with adenovirus vectors containing CA-Rap1a or control LacZ (β-galactosidase) (n= 6 mice/group, mean ± SEM, *p < 0.05, n.s., non-significant).

### Statins Inhibit Rap1a in Isolated Hepatocytes and the Human Liver

Attachment of GGPP to Rap1a’s cysteine residue results in its membrane localization and Epac2-mediated GTP-loading, which supports the importance of the mevalonate pathway in Rap1a’s activation (Goody et al., 2017; Jaskiewicz et al., 2018). Interestingly, a previous report showed that statin treatment of skeletal muscle cells *in vitro* lowers Rap1a isoprenylation, which was suggested as a plausible mechanism to explain statin-mediated myotoxicity (Jaśkiewicz et al., 2018). Because the liver is the primary organ responsible for the metabolism and action of statins, we hypothesized that one mechanism by which statins increase new-onset T2D is their ability to lower GGPP, inhibit Rap1a, and aberrantly activate HGP. As a first step in investigating this hypothesis, we determined whether statin treatment of hepatocytes inhibits Rap1a’s prenylation and membrane localization. We found that treatment of primary mouse hepatocytes with two potent statins in clinical use, simvastatin or rosuvastatin, significantly lowered membrane Rap1a protein, without decreasing the total Rap1a levels (**Figures 3A and 3B).** Further, the decrease in membrane localization of Rap1a was associated with inhibition of its activity, as rosuvastatin treatment lowered GTP-bound, active Rap1a levels (**Figure 3C**). To study the relevance to humans, we obtained human liver specimens from the National Institutes of Health–sponsored Liver Tissue Cell Distribution System. Strikingly, we observed that GTP-bound active Rap1a levels were significantly decreased in the livers of human patients on statin therapy compared to their disease-, age- and sex-matched controls who are not using statins (**Figure 3D**). We obtained similar results from liver samples of another set of patients that are matched for age, sex and BMI (**Figure 3E**). Thus, statin treatment inhibits Rap1a activity in mouse hepatocytes and the human liver.

**Figure 3.**
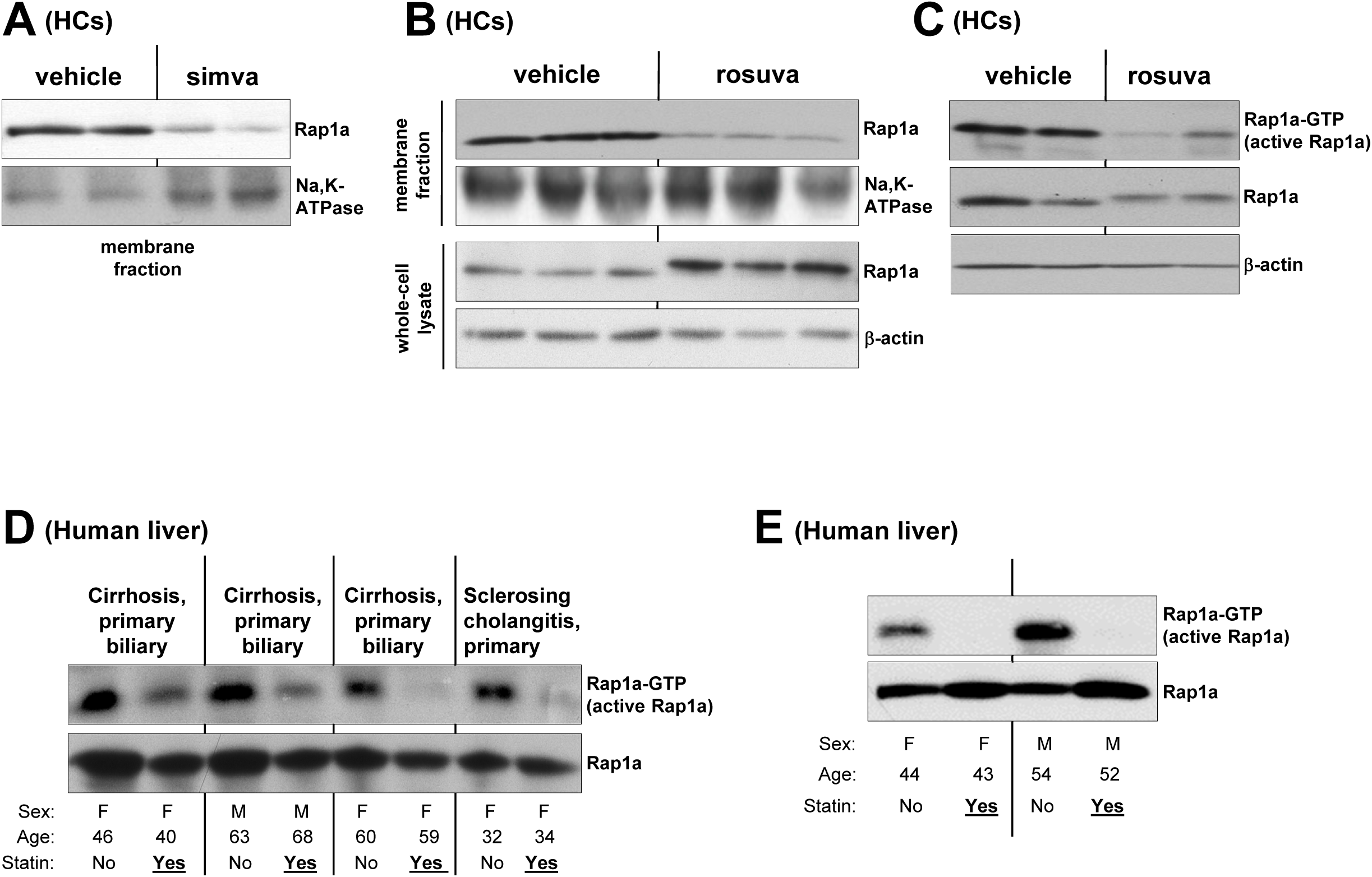
Statins Inhibit Rap1a in Isolated Hepatocytes and the Human Liver. **(A)** Primary mouse hepatocytes (HCs) were treated with vehicle or 10 μM simvastatin (simva) for 20 h. Membrane proteins were assayed for Rap1a and Na, K-ATPase (loading control). **(B)** Same as in (A), except that 5 μM rosuvastatin (rosuva) was used, and membrane fractions and whole-cell lysates were assayed for Rap1a and loading controls (Na, K-ATPase and β-actin, respectively). **(C)** Hepatocytes treated as in (B) were assayed for GTP-bound (active) Rap1a and total Rap1a. **(D)** Liver tissue samples from patients on statin therapy and their sex-, age- and disease-matched controls without statin therapy were assayed for GTP-bound (active) Rap1a and total Rap1a. **(E)** Same as in (D), except that livers from another set of patients were assayed for GTP-bound (active) Rap1a and total Rap1a.

### Statins Increase Gluconeogenesis in WT But Not in Rap1a KO hepatocytes

Given that Rap1a inhibition increases HGP, we then sought to investigate whether an increase in hepatic gluconeogenesis is involved in the diabetogenic effects of statins. We observed that treatment of primary mouse hepatocytes with different statins, including, simvastatin, rosuvastatin, or fluvastatin increased *G6Pc* and *Pck1* mRNAs (**Figures 4A, S3A, S3C and S3D**). Consistent with an increase in gluconeogenic gene expression, glucose production was also increased upon statin treatment (**Figures 4B and S3B**). Moreover, rosuvastatin treatment of metabolism-qualified primary human hepatocytes resulted in an increase in gluconeogenic gene expression, which provides information about the human relevance of our murine hepatocyte studies (**Figures 4C and 4D**). To understand whether inhibition of Rap1a is one of the major mechanisms by which statins induce gluconeogenic gene expression, we incubated control or si-Rap1a–treated hepatocytes with rosuvastatin. Confirming our above results, both silencing Rap1a or rosuvastatin treatment alone increased *G6Pc* mRNA (**Figure 4E**). Importantly, rosuvastatin’s ability to increase *G6Pc* mRNA was abolished in Rap1a-silenced hepatocytes, which supports the notion that inhibition of Rap1a acts in the same pathway as statins (**Figure 4E**). Similar results were obtained in *Pck1* gene expression, suggesting the idea that inhibition of Rap1a is a major mechanism by which statins increase gluconeogenesis (**Figure 4F**). To show relevance *in vivo*, we treated DIO mice with a high-fat diet containing 0.02 % simvastatin (w/w) for 12 weeks and observed that both groups gained similar weight during the treatment (**Figure 4G**). As expected, statin treatment resulted in an increase in the mRNA expression of Srebp2 target genes, including *Hmgcs* and *Hmgcr* in the liver (**Figure S3E**). Similar to hepatic Rap1a silencing, we found that statin-treated obese mice had higher fasting blood glucose, increased gluconeogenic gene expression in the liver and were more glucose intolerant compared to control mice, without alterations in plasma insulin levels (**Figures 4G-4L**).

**Figure 4.**
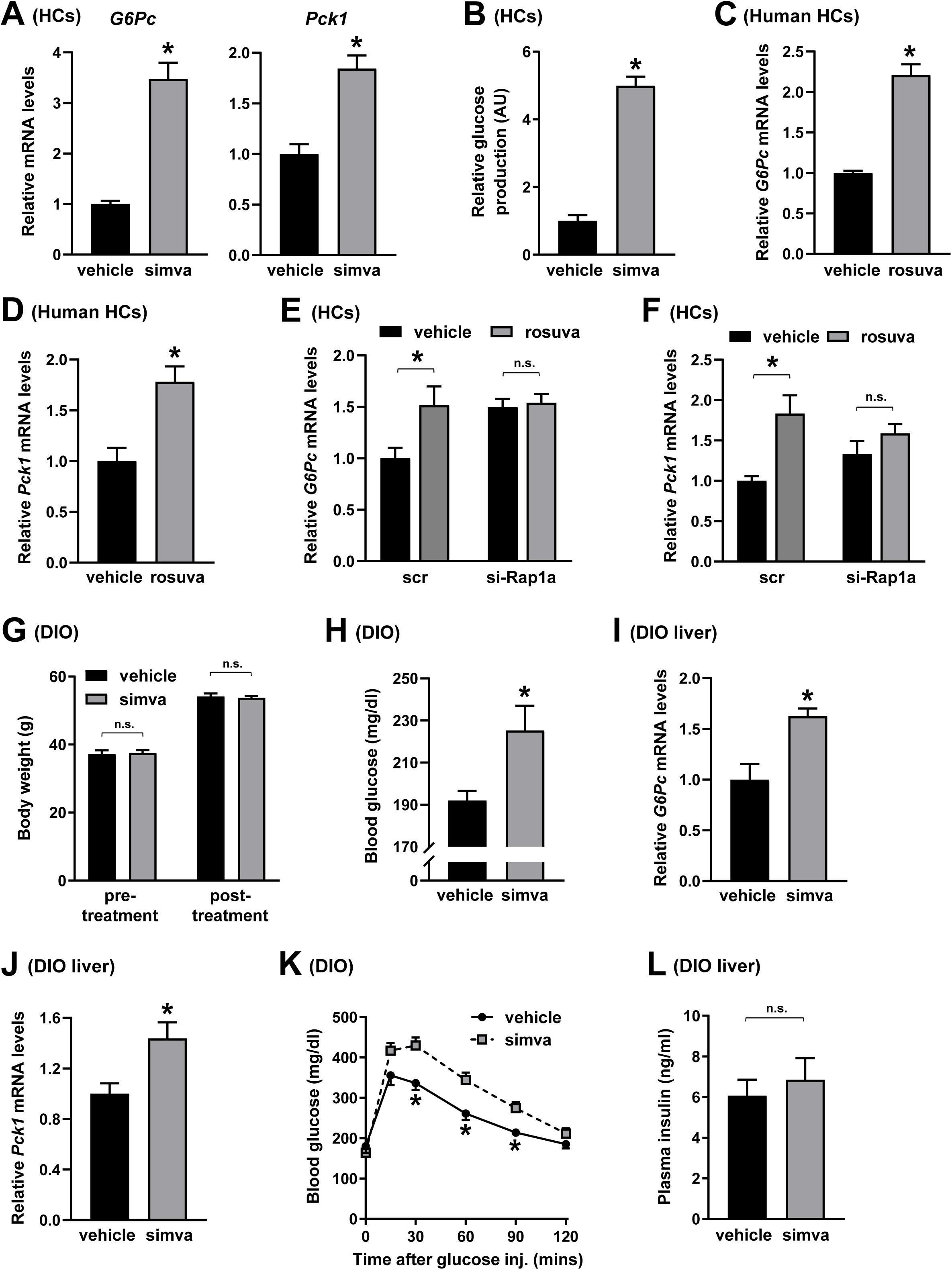
Statins Increase Gluconeogenesis in WT But Not in Rap1a KO Hepatocytes. **(A-B)** Primary mouse hepatocytes (HCs) were treated with vehicle or 10 μM simvastatin (simva) for 24 h and assayed for forskolin and dexamethasone-induced *G6Pc* and *Pck1* mRNA (A) and glucose production (B) (n= 4 wells of cells/group, mean ± SEM, *p < 0.05). **(C-D)** Forskolin and dexamethasone-induced *G6Pc* (C) and *Pck1* (D) mRNAs were measured from primary human hepatocytes treated with vehicle control or 5 μM rosuvastatin (rosuva) for 24 h (n= 4 wells of cells/group, mean ± SEM, *p < 0.05). **(E-F)** Primary mouse hepatocytes that were transfected with scrambled control (scr) or siRNA against Rap1a (si-Rap1a) were treated with vehicle or 5 μM rosuvastatin (rosuva) for 24 h. Forskolin and dexamethasone-induced *G6pc* (E) and *Pck1* mRNA levels (F) were assayed (n= 3 wells of cells/group, mean ± SEM, *p < 0.05, n.s., non-significant). **(G-L)** Body weight before and after the simvastatin treatment (G), 5 h fasting blood glucose (H), liver *G6Pc* (I) and *Pck1* (J) mRNA expression levels, glucose tolerance test (K), and fasting plasma insulin levels (L) from DIO mice that were treated with a high-fat diet containing 0.02 % simvastatin (w/w) (simva) for 12 weeks (n= 6-7 mice/group, mean ± SEM, *p < 0.05, n.s., non-significant).

Statins lower plasma LDL-C via increasing LDLR protein levels and paradoxically increase the expression of PCSK9 (Goldstein and Brown, 2009; Sahebkar et al., 2015; Taylor and Thompson, 2016). To understand the involvement of LDLR and PCSK9 in statin-mediated regulation of hepatocyte glucose metabolism, we treated LDLR and PCSK9 knockout (KO) hepatocytes with rosuvastatin. Similar to WT cells, we observed an increase in *G6Pc* mRNA levels and glucose production in LDLR KO (**Figures S3F and S3G**) and PCSK9 KO hepatocytes treated with rosuvastatin compared to vehicle (**Figures S3H and S3I**). Thus, statins stimulate gluconeogenesis and increase glucose production, and this regulation is independent of LDLR and PCSK9.

### GGPP Supplementation Restores Rap1a Activity and Improves Statin-Induced Glucose Intolerance in Obese Mice

Inhibition of HMG-CoA reductase by statins suppresses GGPP isoprenoid synthesis, which is required for Rap1a to become activated. Because statin treatment or Rap1a inhibition increases gluconeogenesis, we reasoned that a decrease in intracellular GGPP levels is involved in statin-mediated gluconeogenic gene induction. To test this, we put back intracellular GGPP in statin-treated hepatocytes by incubating them with exogenous GGPP. We found that this treatment brought back plasma membrane-localized Rap1a to vehicle-treated levels, suggesting that Rap1a prenylation was restored (**Figure 5A**). Consistent with the restoration of Rap1a’s membrane localization, GGPP treatment abrogated the increase in *G6Pc* and *Pck1* mRNA levels conferred by statin treatment (**Figure 5B**). Of note, GGPP treatment did not inhibit simvastatin induced *Hmgcr* and *Hmgcs* mRNAs (**Figure S4A**), suggesting that induction of Srebp2 by statins was intact and unaffected by GGPP restoration. We next sought to determine if GGPP supplementation in statin-treated obese mice could lower HGP and improve glucose intolerance. Accordingly, we fed DIO mice with 0.02 % simvastatin (w/w)-containing high-fat diet for 12 weeks. Mice were then administered with GGPP precursor, geranylgeraniol (GGOH, 100 mg/kg), or vehicle control by daily gavage while still receiving the statin-containing diet (Gibbs et al., 1999). After 3 weeks of treatment, we observed higher hepatic active-Rap1a levels (**Figure S4B**), lower fasting blood glucose (**Figure 5C),** improved glucose intolerance (**Figure 5D**), and decreased hepatic gluconeogenic gene expression (**Figure 5E**), without a change in body weight or plasma insulin levels (**Figures 5F-5G**) in GGOH-supplemented group as compared to statin treatment alone. Of note, we did not observe changes in blood glucose, hepatic gluconeogenic gene expression, body weight or plasma insulin levels in DIO mice treated with GGOH alone (**Figures S4C-4G**).

**Figure 5.**
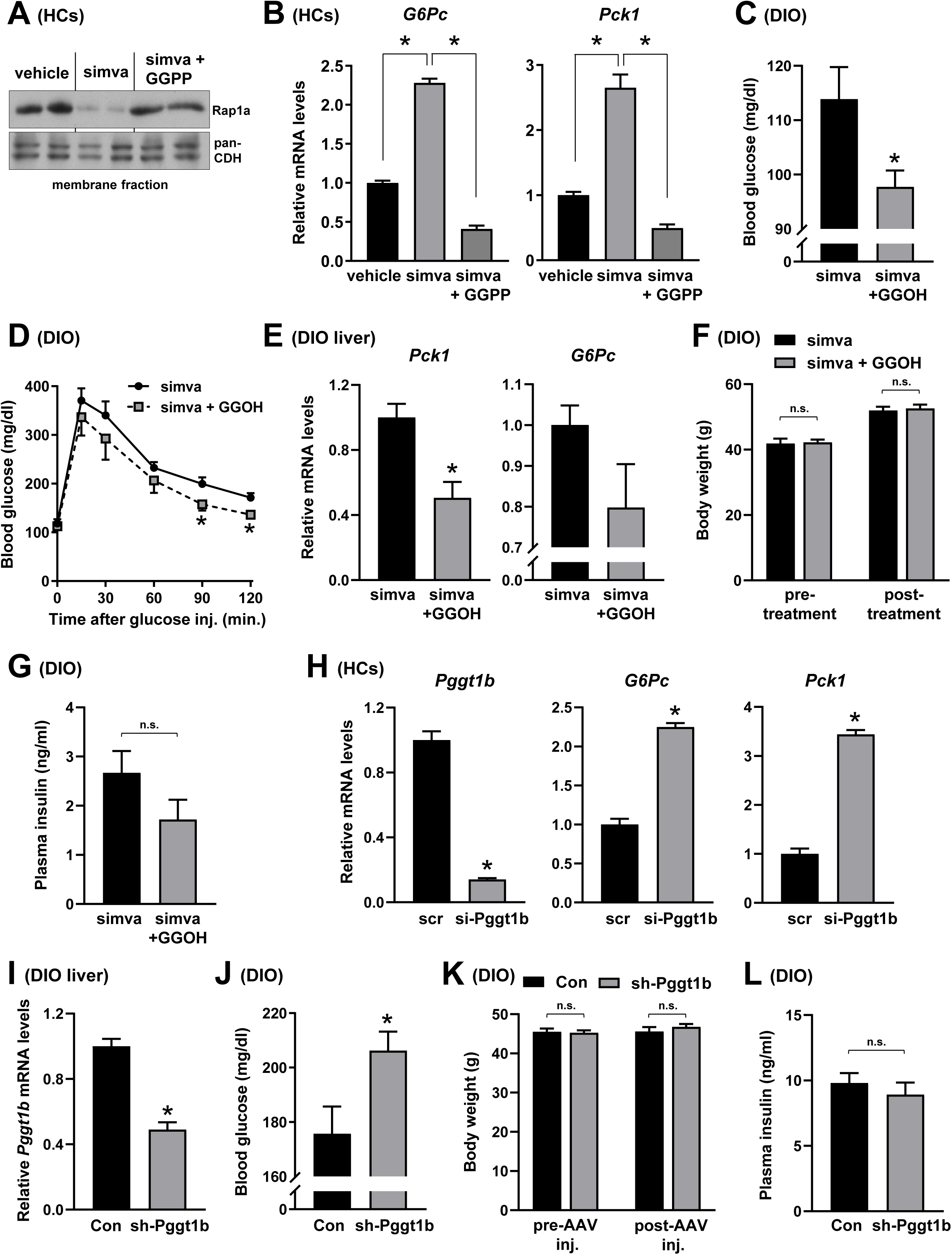
Geranylgeranylation Mediates the Gluconeogenic Effect of Statins. **(A-B)** Primary mouse hepatocytes (HCs) were treated with vehicle, 10 μM simvastatin (simva) or simvastatin + geranylgeranyl pyrophosphate (GGPP) (10 μM) for 24 h. Membrane proteins were assayed for Rap1a and pan-Cadherin (pan-CDH, loading control) (A), and forskolin and dexamethasone-induced *G6Pc* and *Pck1* mRNA levels were measured (B) (n= 3 wells of cells/group, mean ± SEM, *p < 0.05). **(C-G)** DIO mice were fed with 0.02 % simvastatin (w/w)-containing high-fat diet for 12 weeks. Mice were then administered with geranylgeraniol (GGOH, 100 mg/kg/day), or vehicle control by daily gavage for 3 weeks while still receiving the statin-containing diet. Overnight fasting blood glucose (C), glucose tolerance test (D), liver *G6Pc* and *Pck1* mRNA expression levels (E), body weight before and after simvastatin and GGOH treatment (F), and overnight fasting plasma insulin levels (G) were assayed (n= 5-7 mice/group, mean ± SEM, *p < 0.05). **(H)** Primary mouse hepatocytes that were transfected with si-Pggt1b or scrambled control (scr) were assayed for *Pggt1b,* and forskolin and dexamethasone-induced *G6Pc and Pck1* mRNA levels (n=3 wells of cells/group, mean ± SEM, \**P*<0.05, n.s., non-significant). **(I-L)** Liver *Pggt1b* mRNA expression (I), overnight fasting blood glucose levels (J), body weight before and after AAV injection (K) and overnight fasting plasma insulin levels (L) were assayed from DIO mice that were injected with AAV8 vectors containing sh-RNA against Pggt1b (sh-Pggt1b) or empty, control AAV8 (Con) (n= 7 mice/group, mean ± SEM, *p < 0.05, n.s., non-significant).

Given that one of the major effects of statins is to decrease cholesterol biosynthesis, we next asked whether lowering of intracellular cholesterol participates in the gluconeogenic effects of statins. For this purpose, we restored intracellular cholesterol levels in statin-treated hepatocytes by incubating them with cholesterol-enriched phospholipid liposomes (Lipo-Chol) (Wang et al., 2020). As expected, increased expression of cholesterol synthesis genes, including *Hmgcr* and *Hmgcs*, upon statin treatment were lowered back to control levels in cells treated with Lipo-Chol and statin together (**Figures S5A-5B**). However, restoration of intracellular cholesterol was not able to reverse statin induced *G6Pc* and *Pck1* mRNAs (**Figures S5C-5D**). Notably, Lipo-Chol treatment did not affect membrane localization of Rap1a (**Figure S5E**). These results support the hypothesis that statins inhibit Rap1a activity and increase HGP via lowering GGPP levels, which is independent of a change in intracellular cholesterol content.

### Geranylgeranyl Transferase-1 Inhibition Mimics Statins and Increases Gluconeogenesis and HGP

Our results thus far suggest that statins inhibit membrane localization and activation of Rap1a and increase HGP and restoring GGPP levels brings back Rap1a activity and lowers glucose production. Geranylgeranyl transferase 1 (GGT1) covalently attaches GGPP to the cysteine residue in the C-terminal CAAX motif of Rap1a to form a stable thioether bond (Jaskiewicz *et al*., 2018). This post-translational modification increases Rap1a’s hydrophobicity, which results in Rap1a’s plasma membrane localization (Ghomashchi et al., 1995). To understand the contribution of GGT1 to hepatic glucose metabolism, we used a specific GGT1 inhibitor or silenced *Pggt1b*, the gene encoding GGT1, in primary hepatocytes. Similar to statin treatment or Rap1a silencing, inhibition or silencing of GGT1 increased gluconeogenic gene expression and glucose production in both primary mouse and human hepatocytes (**Figures 5H and S6A-S6D**). Consistent with a role for GGT1 in gluconeogenesis *in vivo*, treatment of DIO mice with hepatocyte-specific AAV8 encoding short hairpin RNA (shRNA) construct targeting *Pggt1b* resulted in lowering of hepatic *Pggt1b* mRNA expression by ∼50% and increased fasting blood glucose levels in obese mice, without altering body weight or plasma insulin (**Figures 5I-5L**). It is noteworthy to mention that farnesyl transferase inhibitor FTI-277 did not affect *G6Pc* or *Pck1* mRNA levels in primary mouse hepatocytes (**Figure S6E**). Collectively, these data suggest that inhibition of GGPP synthesis and Rap1a activity by statins increases HGP, which is mimicked by GGT1 inhibition.

### Rap1a Inhibition or Statin Treatment Increases Gluconeogenesis via Regulating FoxO1 Activity

In an attempt to understand the molecular mechanism(s) by which Rap1a regulates HGP, we first determined whether Rap1a deficiency in primary hepatocytes affects intracellular cAMP levels. We found that both basal and forskolin and dexamethasone-treated intracellular cAMP levels were similar in Rap1a-deficient or CA-Rap1a-overexpressing cells compared to control (**Figures S7A-S7B**). Protein kinase A (PKA) is well-known to stimulate gluconeogenesis via increasing the nuclear localization and activity of CREB and CRTC2 transcription factors (Chrivia et al., 1993; Wang et al., 2012b). We next considered the possibility that Rap1a deficiency may stimulate gluconeogenesis via activating PKA. However, we observed that the levels of both p-CREB and p-PKA substrates, as measures of CREB and PKA activities, respectively, were not increased in Rap1a-silenced hepatocytes versus control (**Figure S7C**). In support of this finding, one of the downstream effectors for both CTRC2 and CREB pathways, peroxisome proliferator-activated receptor-γ coactivator-1α (*Pgc1a*) mRNA levels (Herzig et al., 2001) were similar in WT and Rap1a-deficient hepatocytes (**Figure S7D**). Interestingly, *Pgc1a* mRNA levels were decreased, not increased, in cells treated with simvastatin or rosuvastatin compared to controls (**Figure S7E**). These results support the idea that Rap1a regulates gluconeogenesis without an effect on intracellular cAMP levels and independent of PKA.

Forkhead box O1 (FoxO1) is a major gluconeogenic transcription factor that induces the transcription of *G6Pc* and *Pck1* (Pajvani and Accili, 2015). To investigate whether statins or Rap1a inhibition enhance gluconeogenesis via regulating FoxO1, we first measured the mRNA levels of another well-known FoxO1 target gene, *Igfbp1* (Matsumoto et al., 2006). We observed that Rap1a deficiency, statin treatment, or GGT1i treatment increased hepatic *Igfbp1* mRNA levels (**Figures 6A-B, S7F-S7G**). Glucokinase (encoded by *Gck*) is critical for hepatic glucose utilization and is negatively regulated by FoxOs (Haeusler et al., 2014; Langlet et al., 2017b). In support of the role of Rap1a in regulating FoxO1, we found lower *Gck* mRNA levels in Rap1a-deficient DIO liver as compared to control (**Figure 6C**). Likewise, *Gck* mRNA was significantly lower in Rap1a KO hepatocytes under both basal and in response to a combination of forskolin, dexamethasone and insulin treatment, which has been shown to induce *Gck* (Langlet et al., 2017a) (**Figure S7H**). Conversely, CA-Rap1a overexpression increased both basal and forskolin + dexamethasone + insulin─induced *Gck* mRNA in WT hepatocytes (**Figure S7I)**. We then addressed whether Rap1a affects FoxO1’s transcriptional activity using reporter assays. For this purpose, we transfected WT or Rap1a KO hepatocytes with control or GFP-FoxO1 plasmids (**Figures 6D and 6E**, lower blots). We observed that the increase in *Igfbp1* and *G6Pc* promoter activities upon FoxO1 overexpression were further upregulated in Rap1a-deficient hepatocytes (**Figures 6D and 6E**). We found similar results upon forskolin and dexamethasone treatment, suggesting that FoxO1’s transcriptional activity is increased in Rap1a-deficient cells (**Figure S7J**). To test the role of FoxO1 in Rap1a- and statin-mediated gluconeogenesis regulation, we silenced Rap1a in FoxO1,3,4 deficient hepatocytes. The data showed that the ability of Rap1a deficiency to increase *G6Pc* and *Igfbp1* mRNAs was abolished in FoxO1,3,4 deficient hepatocytes supporting the involvement of FoxO proteins in Rap1a-mediated gluconeogenic gene regulation (**Figure 6F**). Because nuclear localization of FoxO1 is required for its functional activity, we next measured nuclear FoxO1 levels in Rap1a-deficient or statin-treated hepatocytes. We observed that both Rap1a silencing and statin treatment increased nuclear FoxO1 without altering its total cellular levels (**Figures 6G and 6H**). One of the major upstream regulators of FoxO1’s subcellular localization is insulin receptor signaling-activated serine/threonine kinase, Akt. Activation of Akt phosphorylates FoxO1, which results in FoxO1’s cytoplasmic retention (Brunet et al., 1999). We then sought to determine if phosphorylation of FoxO1 or Akt was altered upon Rap1a deficiency. We found that Rap1a-silenced hepatocytes had significantly lower phospho-Akt and phospho-FoxO1 levels upon insulin stimulation or under basal conditions (**Figures 6I and 6K**). Similarly, rosuvastatin treatment of hepatocytes impaired insulin induced Akt and FoxO1 phosphorylation (**Figures 6J, 6L, 6M**). These results support the idea that statins or Rap1a inhibition reduce phosphorylation of Akt, which results in nuclear localization of FoxO1 and stimulation of gluconeogenesis.

**Figure 6.**
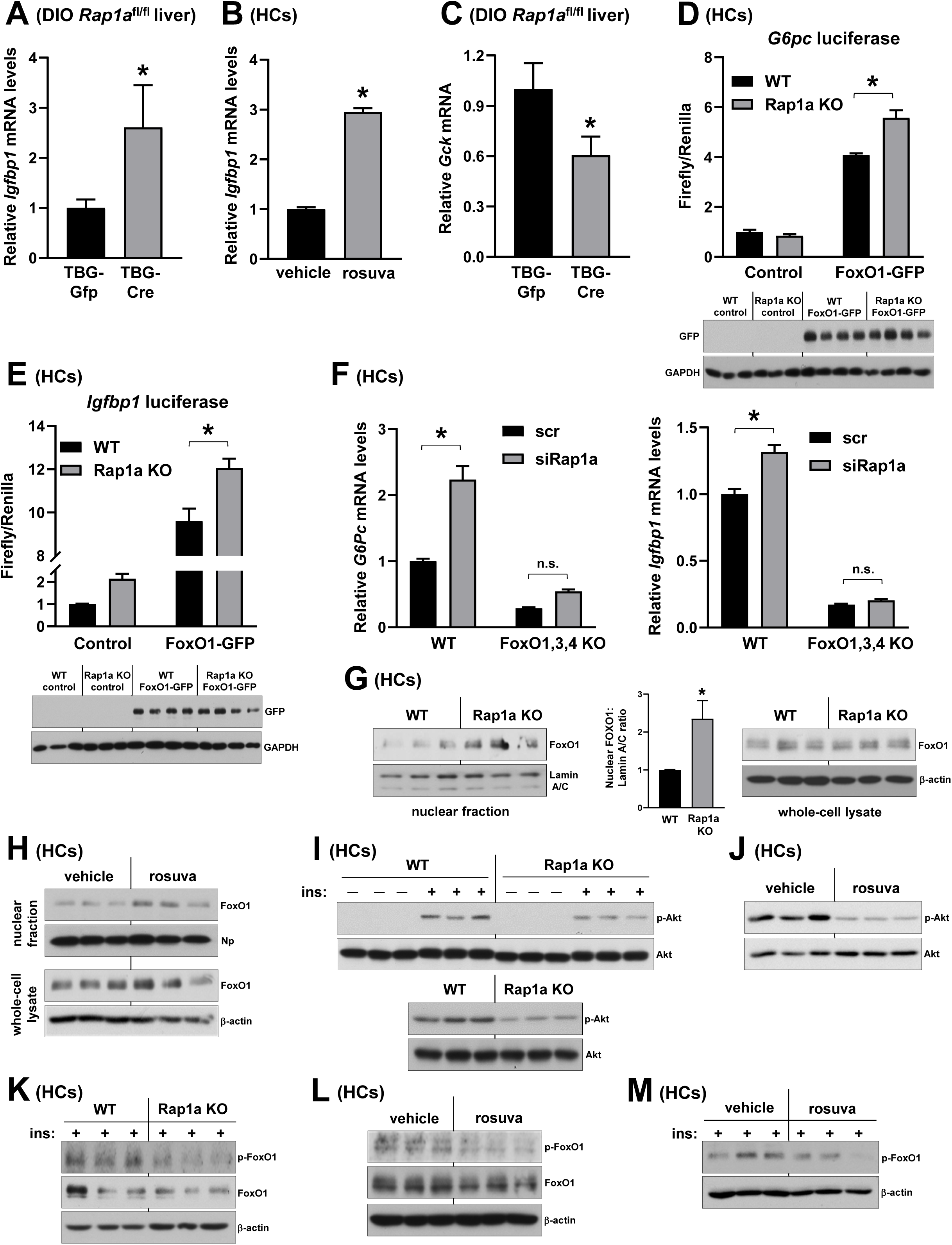
Rap1a Inhibition or Statin Treatment Increases Gluconeogenesis via Regulating FoxO1 Activity. **(A-B)** *Igfbp1* mRNA levels were measured from the livers of DIO *Rap1a*^fl/fl^ mice that were injected with TBG-Cre or TBG-Gfp (A), and vehicle- or 5 μM rosuvastatin (rosuva)-treated primary hepatocytes (HCs) that were incubated with forskolin and dexamethasone (B) (n= 7-8 mice/group and n= 3 wells of cells/group, respectively, mean ± SEM, *p < 0.05). **(C)** *Gck* mRNA levels were measured from the livers of DIO *Rap1a*^fl/fl^ mice that were injected with TBG-Cre or TBG-Gfp (n= 7-8 mice/group, mean ± SEM, *p < 0.05). **(D)** WT and Rap1a KO primary hepatocytes were transfected with a luciferase fusion construct encoding nucleotides −1227 to +57 of the *G6pc* promoter containing an intact FoxO binding site and FoxO1-GFP or control plasmid. Relative luciferase activity and GFP levels from WT and Rap1a KO cells transfected with FoxO1-GFP were measured (lower blots) (n= 3-4 wells of cells/group, mean ± SEM, *p < 0.05). **(E)** Luciferase reporter assay of the *Igfbp1* promoter in WT and Rap1a KO cells transfected with control or FoxO1-GFP. GFP levels from WT and Rap1a KO cells transfected with FoxO1-GFP were assayed in the lower blots (n= 3-4 wells of cells/group, mean ± SEM, *p < 0.05). **(F)** *G6Pc* and *Igfbp1* mRNA levels from forskolin and dexamethasone-treated WT and FoxO1,3,4 KO cells that were transfected with scrambled RNA (scr) or siRNA against Rap1a (si-Rap1a) (n= 4 wells of cells/group, mean ± SEM, *p < 0.05, n.s., non-significant). **(G)** Nuclear and whole-cell FoxO1 levels along with loading controls (Lamin A/C and β-actin, respectively) in WT versus Rap1a KO hepatocytes. Densitometric quantification of the nuclear FoxO1 immunoblot data is shown in the bar graph. **(H)** Same as in (G) except that vehicle or 5 μM rosuvastatin (rosuva)-treated cells were used. **(I)** WT and Rap1a KO hepatocytes were stimulated with vehicle control or insulin (100 nM) for 5 minutes, and phospho-Akt and total Akt levels were assayed (upper blots). Basal levels of phospho-Akt and total Akt in WT and Rap1a KO cells are shown in the lower blots. **(J)** Same as in (I) except that vehicle or 5 μM rosuvastatin-treated cells were used. **(K)** p-FoxO1, total FoxO1 and β-actin levels were measured from WT versus Rap1a KO hepatocytes stimulated with insulin (100 nM) for 5 minutes. **(L-M)** Same as in (K) except that vehicle or 5 μM rosuvastatin-treated cells without (L) or with insulin stimulation (M) were used.

### Rap1a Induces Actin Polymerization and Actin Remodeling Contributes to Gluconeogenesis

We next sought to investigate possible mechanisms linking Rap1a activation to Akt-mediated FoxO1 regulation. Previous work in other cell types have demonstrated that Rap1a promotes filamentous actin (F-actin) polymerization (Mun and Jeon, 2012; Wang et al., 2017). As disruption of actin polymerization and cytoskeleton remodeling inhibits insulin induced Akt activation (Lee et al., 2013; Peyrollier et al., 2000; Wang et al., 2012a), we considered the hypothesis that Rap1a may regulate gluconeogenesis through its effects on actin organization. We first examined whether Rap1a plays a role in actin polymerization in hepatocytes by staining F-actin filaments with Alexa Fluor 555–conjugated phalloidin in scrambled control or si-Rap1aLtreated cells. We observed that si-Rap1aLtreated hepatocytes had disrupted cytoskeletal structures as evidenced by short, fragmented F-actin filaments that coalesce to form cytoskeletal clumps (**Figure 7A,** see white arrows). This fragmented actin phenotype was similar to the actin phenotype of WT cells treated with the potent actin polymerization inhibitor, Cytochalasin D (**Figure 7A**), suggesting that Rap1a silencing results in dysregulation of actin cytoskeleton in hepatocytes. We next asked whether Rap1a inhibition induces gluconeogenesis via blocking actin polymerization. Consistent with this idea, we found that treatment of hepatocytes with different concentrations of Cytochalasin D increased *G6Pc* and *Pck1* mRNAs and resulted in a small but significant increase in glucose production (**Figures 7B and 7C**). We observed similar results when actin polymerization was inhibited using another actin polymerization inhibitor, Latrunculin A (not shown). Conversely, treatment of primary hepatocytes with the actin filament polymerizing and stabilizing agent, Jasplakinolide, lowered gluconeogenic gene expression and glucose production (**Figures 7D and 7E**). Similar to Rap1a inhibition, Cytochalasin D was unable to induce *G6Pc* and *Igfbp1* mRNAs in FoxO1,3,4 deficient hepatocytes (**Figure 7F**). Most importantly, nuclear FoxO1 levels were increased whereas insulin-stimulated phospho-Akt levels were decreased in Cytochalasin D-treated hepatocytes (**Figures 7G and 7H**). Taken together, these data support the idea that hepatocyte Rap1a is involved in actin cytoskeleton dynamics, which contributes to the regulation of gluconeogenesis via Akt-mediated FoxO1 activation.

**Figure 7.**
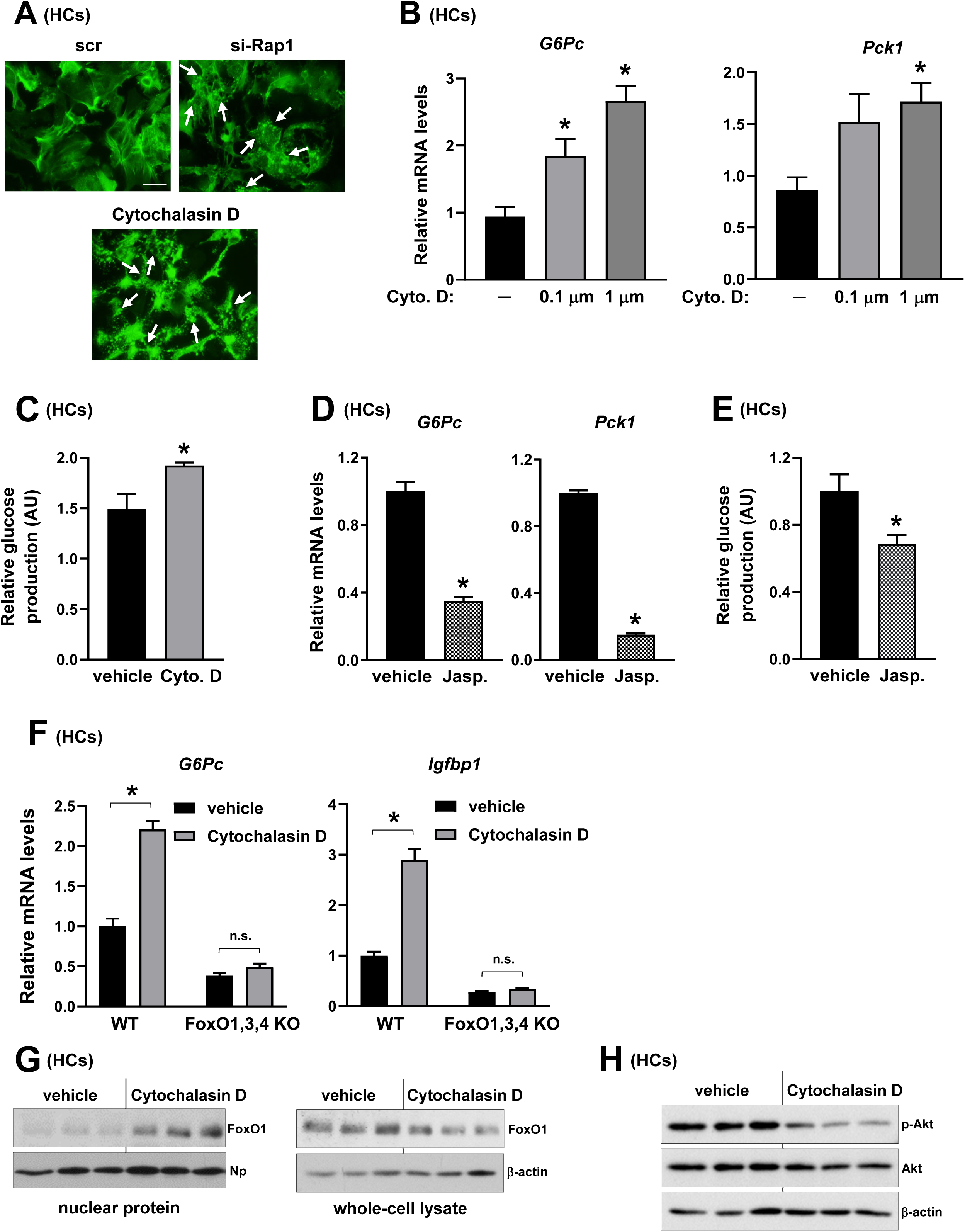
Actin Remodeling Contributes to Gluconeogenesis and Hepatocyte Glucose Production. **(A)** Primary mouse hepatocytes (HCs) that were treated with scrambled control (scr), si-Rap1a or Cytochalasin D were stained with 488-phalloidin (green) to visualize F-actin. White arrows indicate fragmented F-actin filaments that form cytoskeletal clumps. Scale bar 10 μm. **(B)** Hepatocytes were treated with vehicle, 0.1 μM, or 1 μM Cytochalasin D (Cyto. D) for 8 h, and forskolin and dexamethasone stimulated *G6pc* and *Pck1* mRNA levels were measured (n= 4 wells of cells/group, mean ± SEM, *p < 0.05). **(C)** Same as in (B) except that glucose production was measured in cells treated with vehicle or 0.1 μM Cytochalasin D (Cyto. D) (n= 3 wells of cells/group, mean ± SEM, *p < 0.05). **(D)** Hepatocytes were treated with 0.1 μM Jasplakinolide (Jasp.) for 6 h and forskolin and dexamethasone stimulated *G6pc* and *Pck1* mRNA levels were measured (n= 3 wells of cells/group, mean ± SEM, *p < 0.05). **(E)** Same as in (D), except that glucose production was measured (n= 4 wells of cells/group, mean ± SEM, *p < 0.05). **(F)** WT and FoxO1,3,4 KO cells were treated with vehicle or 1 μM Cytochalasin D for 8 h, and forskolin and dexamethasone stimulated *G6pc* and *Igfbp1* mRNA levels were measured (n= 4 wells of cells/group, mean ± SEM, *p < 0.05, n.s., non-significant). **(G)** Nuclear and whole-cell FoxO1 levels along with loading controls (nucleophosmin, Np, and β-actin, respectively) in hepatocytes that were treated with vehicle or 1 μM Cytochalasin D for 8 h. **(H)** Same as in (G), except that cells were stimulated with insulin (100 nM) for 5 minutes, and phospho-Akt, total Akt and β-actin levels were assayed.

## DISCUSSION

The combination of insulin resistance together with unopposed glucagon action results in increased HGP, which is largely responsible for the fasting hyperglycemia observed in obese patients with T2D (Gastaldelli et al., 2000; Unger and Cherrington, 2012). The most commonly used anti-diabetes therapy, metformin, is well-known for its ability to decrease hepatic gluconeogenesis, and other candidates that reduce HGP are being investigated as anti-glycemic agents (Kazda et al., 2016; Lee et al., 2021; Sharabi et al., 2017; Vella et al., 2019). Our results here identify the small GTPase, Rap1a, as a new potential target that regulates HGP and glucose homeostasis. Mechanistically, we show that Rap1a stimulates actin polymerization, which regulates FoxO1’s nuclear localization and activity. The role of FoxO1 in Rap1a action is supported by the finding that the stimulatory effects of inhibition of Rap1a or actin polymerization on gluconeogenesis are abrogated in FoxO1,3,4 deficient hepatocytes. With regard to FoxO1 regulation, we provide data that phosphorylation and activation of Akt, the major kinase that inhibits FoxO1 activity, is suppressed upon Rap1a-deficieny. Of note, previous studies have implicated actin network and Rap1 in Akt activation through their effects on phosphoinositide 3-kinase (PI3K) activity, the upstream Akt regulator (Cahill et al., 2016; Eyster et al., 2005; Kortholt et al., 2010). Because of the importance of insulin receptor-Akt pathway in the pathogenesis of many obesity-associated metabolic dysfunctions, future investigations will be necessary to determine the molecular mechanisms linking Rap1a to Akt activation in hepatocytes.

In addition to obesity-induced hyperglycemia, our results show that inhibition of Rap1a activity or GGPP synthesis mediates elevated HGP in statin-treated hepatocytes and mice, which could be one of the underlying mechanisms linking statins to new-onset T2D. Interestingly, a recent work from another group independently reported that restoring geranylgeranyl isoprenoids improves statin associated glucose intolerance (Wang et al., 2022). *In vitro* findings from Wang et al. suggested that geranylgeranylation of another GTPase, Rab14, contributes to hepatic insulin signaling; however, the importance of Rab14 in statin associated glucose intolerance was not studied. Given that rosuvastatin’s ability to increase gluconeogenic genes was abolished in Rap1a deficient hepatocytes (Figures 4E and 4F), our results suggest that inhibition of Rap1a is one of the major underlying mechanisms by which statins increase gluconeogenesis.

Because all statins have high selectivity for the liver, largely because of efficient first-pass uptake (Schachter, 2005), the liver likely plays a major role in statin-induced glucose intolerance. However, statins may also contribute to glucose intolerance through their effects on other insulin-sensitive tissues and the pancreas. For example, statin treatment was shown to inhibit insulin secretion in pancreatic β-cell lines and DIO mice. (Henriksbo et al., 2019; Hwang et al., 2019; Salunkhe et al., 2016). Interestingly, a recent study in human subjects treated with atorvastatin for 10 weeks reported an increase in steady-state plasma glucose without a decline in insulin secretion (Abbasi et al., 2021), yet the effect of long-term statin-use on β-cell function remains to be investigated. It is important to note that activation of Epac2 and Rap1 in β-cells is suggested to be beneficial in glucose homeostasis, as Epac2─Rap1 activation results in increased incretin-mediated insulin secretion (Takahashi et al., 2015). Regarding other possible non-hepatic effects, previous work showed impairment in insulin signaling in cultured adipocytes and myocytes upon statin treatment. Moreover, a recent study reported that GGPP and GGT1 are important regulators of brown adipocyte function and contribute to systemic glucose metabolism (Balaz et al., 2019). Future *in vivo* studies will be needed to determine whether geranylgeranylation or Rap1a in different cell types contribute to the statin-T2D link.

Clinical trials of PCSK9 inhibitors have not shown an effect of these drugs on glycemia or T2D diagnosis; however, an increased T2D risk has been observed in individuals with PCSK9 loss-of-function variants (Colhoun et al., 2016; Ference *et al*., 2016; Lotta et al., 2016; Sabatine et al., 2017; Sattar et al., 2017; Schmidt et al., 2017). On the other hand, LDL-C–modulating allele in *APOB* showed no relationship with diabetes risk (Xu et al., 2017). Thus, T2D link with statins and PCSK9 loss-of-function variants is complex and may not be simply attributed to their effects on liver LDLR expression or plasma LDL-C, which is in line with our data showing that intracellular cholesterol or LDLR do not impact statin-induced gluconeogenesis in hepatocytes. Finally, while our results have defined an important mechanism that may underlie statin’s hyperglycemic effects, it is important to note that the benefits of statin-use for the reduction of cardiovascular disease risk outweigh the risk of developing T2D in the majority of patients, especially in individuals with higher cardiovascular disease risk (Mach et al., 2018; Sattar et al., 2014).

In sum, our previous work demonstrated that hepatic Rap1a activation contributes to plasma LDL-C regulation via lowering PCSK9 (Spolitu *et al*., 2019). The data present here show that Rap1a regulates HGP, and blocking Rap1a prenylation, through lowering intracellular GGPP isoprenoid levels, contributes to statin-induced glucose intolerance. These results suggest new roles for Rap1a and provide new insight into the metabolic regulation.

## ACKNOWLEDGMENTS

We thank Dr. Rebecca Haeusler (Columbia University) for *Foxo1,3,4*^fl/fl^ mice, and Dr. Keith Burridge and Dr. Erika Wittchen (University of North Carolina at Chapel Hill) for CA-Rap1a adenovirus. We would like to acknowledge the Gene Therapy Resource Program (GTRP) of the NHLBI for providing AAV8-TBG-Cre and AAV8-TBG-Gfp, and NIH-supported Liver Tissue Cell Distribution System at the University of Minnesota for arranging and providing de-identified human liver samples (NIH Contract #HSN276201200017C). This work was supported by State Scholarship Fund from China Scholarship Council (201906370237) to Y.W.; Medical Student Research Training Grant (2T32DK007559-27A1) to J.A.Z.; NIH grant (DK124457) and Russell Berrie Foundation Pre-translational Diabetes Research Award to L.O.

## AUTHOR CONTRIBUTIONS

Y.W., S.S., J.A.Z and A.S. designed and performed the experiments, analyzed the data, interpreted results, and edited the manuscript. L.O. conceived and supervised the project, designed experiments, analyzed data, interpreted results, and wrote the manuscript.

## STAR METHODS

### Mouse experiments

WT C57BL/6J (stock number: 000664), *db/db* and *db/+* (stock number: 000642) and diet-induced obese (DIO, stock number: 380050) mice and their controls (stock number: 380056) were from Jackson Labs. *Foxo1,3,4*^fl/fl^ mice were generously provided by Dr. Rebecca Haeusler from Columbia University. Floxed *Rap1a* mice (Jackson Labs, stock number: 021066) were generated by crossbreeding *Rap1ab* double floxed mice with WT C57BL/6J mice and confirmed by genotyping. 4-week-old mice were fed a high-fat diet with 60% kcal from fat (Research Diets, D12492) for 12-20 weeks. To obtain hepatocyte-specific *Rap1a* knockout mice, 16-20-week-old *Rap1a*^fl/fl^ mice were injected with adeno-associated virus (AAV) expressing Cre recombinase, driven by the thyroxin-binding globulin (TBG) promoter (AAV8-TBG-Cre). Littermates injected with AAV8-TBG-Gfp were used as control mice. Recombinant AAVs (1–2.5 × 10^11^ genome copy per mouse) and adenoviruses (0.5–0.75 × 10^9^ plaque-forming units per mouse) were delivered by tail vein injections. For statin treatment experiments, the mice were fed a high-fat diet containing 0.02% (w/w) simvastatin (Research Diets, D12492) for 12 weeks. Geranylgeraniol (GGOH) was given by oral gavage at a dose of 100 mg/kg/day for 3 weeks. Fasting blood glucose was measured in mice that were fasted for 5 h or overnight, with free access to water, using a glucose meter (OneTouch Ultra). Plasma insulin levels were measured in mice that were fasted for 5 h using ultrasensitive mouse insulin ELISA Kit (Crystal Chem, cat # 90080). Glucose tolerance tests (GTT) were performed in overnight-fasted mice by assaying blood glucose at various times after intraperitoneal injection of glucose (0.5 g/kg for *db/db* mice and 1.0-1.5 g/kg for DIO mice). All mice were maintained on a 12-12-h light-dark cycle. For all experiments, male mice of the same age and similar weight were randomly assigned to experimental and control groups. Animal studies were performed in accordance with the Columbia University Institutional Animal Care and Use Committee.

### Human samples

Human liver samples were obtained from the NIH-supported Liver Tissue Cell Distribution System at the University of Minnesota. The samples were collected postmortem on the date of liver transplantation and preserved as frozen samples. Part of the human liver samples were obtained from patients undergoing bariatric surgery or clinically indicated laparoscopic procedures at the New York Presbyterian Hospital, Columbia University Irving Medical Center. Samples were obtained from intra-operative needle biopsies of the liver at a standard anatomic location. The biopsy specimens were frozen immediately in liquid nitrogen and stored at −80°C until subsequent analyses. The Institutional Review Board at the Columbia University Medical Center approved the research protocol. All participants provided written informed consent.

### Reagents and antibodies

Forskolin (cat # F6886), dexamethasone (cat # D4902), GGT1i (cat # G5294 and G5169), FTi (cat # F9803), Fluvastatin (cat# SML0038), glucose (cat #G7021), glucagon (cat # G2044) and insulin (cat # I0516) were from Sigma. Simvastatin for cell treatment was from Sigma (cat # S6196) and simvastatin for mouse experiments was from Tokyo Chemical Industry (TCI) Company (cat # S0509). Rosuvastatin (cat # 18813), GGPP (cat # 63330) and GGOH (cat # 13272) were from Cayman Chemicals. Anti-β-actin (cat # 4970), anti-pan-Cadherin (cat # 4068), anti-GFP (cat # 2956), anti-GAPDH (cat # 5174), anti-p-Akt (cat # 4060), anti-Akt (cat# 4691), anti-FoxO1 (cat # 2880), anti-p-FoxO1 (cat# 9464), anti-Lamin A/C (cat # 4777), anti-nucleophosmin (anti-NPM) (cat # 3542), anti-p-CREB (cat # 9198), anti-CREB (cat # 9197) and anti-p-PKA substrates (cat # 9624) antibodies were from Cell Signaling Technology. Anti-Rap1a antibody (cat # AF3767) was from R&D Systems. Anti-Na,K-ATPase antibody (cat # ab76020) was from Abcam. siRNAs were purchased from Integrated DNA Technologies and designed as following: siEpac2, 5’- rArArGrCrArArCrArGrArUrUrCrGrGrUrUrUrUrArArArUGA-3’ (sense), 5’- rUrCrArUrUrUrArArArArCrCrGrArArUrCrUrGrUrUrGrCrUrUrCrA-3’ (antisense), siRap1a, 5’-rCrArArGrCrUrArGrUrArGrUrCrCrUrUrGrGrUrUrCrArGGA-3’ (sense), 5’- rUrCrCrUrGrArArCrCrArArGrGrArCrUrArCrUrArGrCrUrUrGrUrA-3’ (antisense), siPggt1b, 5’-rCrUrUrArArGrGrUrGrUrGrCrCrArArCrUrArArArCrArUGT-3’ (sense), 5’- rArCrArUrGrUrUrUrArGrUrUrGrGrCrArCrArCrCrUrUrArArGrArA-3’ (antisense). AAV8-TBG-Cre and AAV8-TBG-Gfp were obtained from Gene Therapy Resource Program (GTRP) of NHLBI or purchased from Addgene. AAV8-shRNA targeting murine Pggt1b was made by annealing complementary oligonucleotides and then ligating them into the pAAV-RSV-GFP-H1 vector, as described previously (Lisowski et al., 2014). The resultant constructs were amplified by Salk Institute Gene Transfer, Targeting, and Therapeutics Core. AAV8-TBG-CA-Rap1a was from Penn Vector Core. All plasmids were amplified by Genewiz. Adenoviruses encoding LacZ and CA-Rap1a were described previously (Wittchen *et al*., 2011) and amplified by Viraquest, Inc. (North Liberty, IA).

### Hepatocyte experiments

Primary mouse hepatocytes were isolated from 8- to 14-week-old male and female mice as described previously (Ozcan et al., 2012). AML-12 cells were from ATCC (Cat # CRL-2254). Hepatocytes were cultured in DMEM/F-12 containing 10% FBS (fetal bovine serum) and treated as described in the figure legends. Transfections with siRNA or plasmids were carried out using Lipofectamine RNAiMAX or Lipofectamine 2000 reagents according to manufacturer’s instructions. For gluconeogenesis, cells were cultured with DMEM/F-12 containing 0% FBS for the last three hours and stimulated with 10 μM forskolin plus 100 nM dexamethasone (F+D) or 100 nM glucagon. For glucose production assays, primary hepatocytes were cultured in regular DMEM-F12 medium. Cells were then washed three times with warmed PBS and then incubated with glucose-free DMEM-F12 without phenol red media containing 2 mM sodium pyruvate, 20 mM lactate, 100 nM dexamethasone, and 10 μM forskolin for sixteen hours. Culture media was collected for glucose concentration measurement using Glucose (GO) Assay Kit (Sigma, cat # GAGO20), and the cells were washed with PBS and lysed in RIPA buffer for protein determination. Glucose production was normalized to the protein amount of the cells and presented as relative to control.

### Immunoblotting

Total liver or cell protein was lysed using RIPA buffer (Thermo Scientific) containing 2 mM PMSF, 5 µg/ml leupeptin, 10 nM okadaic acid and 5 µg/ml aprotinin on ice. Protein concentration was measured by a BCA assay kit (Bio-Rad). Protein extracts were electrophoresed on SDS-polyacrylamide gels and transferred to 0.45 µm or 0.2 µm PVDF membranes. Blots were blocked in Tris-buffered saline with 0.1% Tween-20 containing 5% BSA at room temperature for one hour. Membranes were then incubated overnight at 4°C with primary antibodies. The protein bands were detected with horseradish peroxidase-conjugated secondary antibodies (Jackson Immuno Research) and Supersignal West Pico enhanced chemiluminescent solution (Thermo). ImageJ was used for densitometric analysis of the immunoblots.

### Membrane protein extraction

Membrane proteins were isolated as described previously (Petersen et al., 2016). Briefly, sells were washed with cold HBSS and lysed with buffer A containing 20 mM Tris-HCl pH7.4, 1 mM EDTA, 0.25 mM EGTA, 250 mM Sucrose, 10 mM NaF, and 2 mM Na_3_VO_4_ on ice. Cell lysates were collected and centrifuged at 500 x g for 10 minutes at 4°C. The supernatant was collected and centrifuged at 100,000 x rpm for 1 hour at 4°C. Supernatant was discarded, and the pellet was re-suspended with buffer B containing 250 mM Tri-HCl pH7.4, 1 mM EDTA, 0.25 mM EGTA, 10 mM NaF, 2 mM Na_3_VO_4_ and 2% Triton-X on ice for 30 minutes. Lysates were centrifuged at 15,000 x rpm for 20 minutes at 4°C. Supernatant was collected, and protein concentration was measured.

### Nuclear protein extraction

A modified Active Motif Nuclear Kit 40010 was used for nuclear protein extraction. Cells were scraped with cold PBS containing 2 mM PMSF, 5 µg/ml leupeptin, 10 nM okadaic acid and centrifuged at 200 x g for 5 minutes at 4°C. Supernatant was discarded, and the pellet was re-suspended with 1X hypotonic buffer and incubated on ice for 15 minutes followed by detergent treatment. The lysates were then centrifuged at 14,000 x g for 30 seconds at 4°C. Supernatant was discarded, and the pellet was re-suspended in RIPA buffer containing 2 mm PMSF, 5 µg/ml leupeptin, 10 nM okadaic acid and 5 µg/ml aprotinin for 30 minutes on ice. Lysates were centrifuged at 14,000 x g for 10 minutes at 4°C, and the supernatant was collected, and protein concentration was measured.

### Rap1a activity

Rap1a activity was assayed using Active Rap1 Detection Kit (Cell Signaling Technology). Cells were washed with cold PBS and lysed with lysis/binding/washing buffer containing 2 mM PMSF, 5 µg/ml leupeptin, 10 nM okadaic acid and 5 µg/ml aprotinin. GST-RalGDS-RBD fusion protein was used to bind the activated form of GTP-bound Rap1, which was then immunoprecipitated with glutathione resin. Rap1a activation levels were determined by western blot using a Rap1a primary antibody.

### Measurement of cAMP

Cyclic AMP XP^®^ Assay Kit (Cell Signaling Technology) was used for cAMP measurements. Cells were lysed with lysis buffer and centrifuged briefly to remove cell debris. Clear lysates were transferred to the cAMP rabbit monoclonal antibody-coated microwells, mixed with HRP-linked cAMP solution, and incubated at room temperature for 3 hours. Following washing to remove excess sample cAMP and HRP-linked cAMP, HRP substrate TMB is added to develop color and then absorbance was measured.

### Transcriptional reporter assays

Hepatocytes were transfected with Igfbp1-promoter/pGL3 (Addgene, cat # 12146) and GFP-FoxO1 plasmids (Addgene, cat # 17551), and renilla luciferase vector was co-transfected as an internal control (Addgene, cat # 27163). Cells were collected 72 hours post-transfection with 1X Passive Lysis Buffer (Promega). Firefly luciferase reporter was determined by addition of Luciferase Assay Substrate and quantification of luminescence on a FLUOstar Omega plate reader.

### Preparation of cholesterol rich liposomes

DMPC (1,2-dimyristoyl-sn-glycero-3-phosphocholine, Avanti Polar Lipids, cat # 850345) and cholesterol (Sigma, cat # C8667) were dissolved in chloroform. For cholesterol rich liposomes, 40 mg of DMPC was mixed with 80 mg of cholesterol in a glass vessel and the solvent was dried under a stream of nitrogen gas. Multilamellar vesicles (large liposomes>1uM) were formed by addition of 10 ml of PBS to the lipid film and then shaking the mixture. The liposomes were reduced in size by probe sonication on ice for 10-15 minutes. The solution turned white and was centrifuged at 10,000 x g for 10 minutes. Supernatant was collected and extruded through 100 nm polycarbonate filter (Avanti, 610000-1Ea). Aliquots were stored in glass vials under argon at 4°C and used within 2 weeks.

### Phalloidin staining

Hepatocytes cultured in glass coverslips were washed with PBS and fixed with 4% formaldehyde for 20 minutes at room temperature. The cells were then permeabilized with 0.1% Triton X-100 in PBS for 5 minutes, followed by 3 washes of PBS. F-actin fibers were stained for 1 h at room temperature with Phalloidin-iFluor 488 reagent (Abcam, cat# ab176753, 1:1000 diluted in 1%BSA). After washing three times with PBS, images were taken using a Leica epifluorescence microscope (DMI6000B).

### Quantitative PCR

RNA was extracted from hepatocytes or liver tissue using TRIzol (Invitrogen) and cDNA was synthesized from 1u g of RNA using a cDNA synthesis kit (Invitrogen). Real-time PCR was performed using a 7500 Real-Time PCR system and SYBR Green reagents (Applied Biosystems). Table 1 listed the specific primer sets.

**Table 1.**
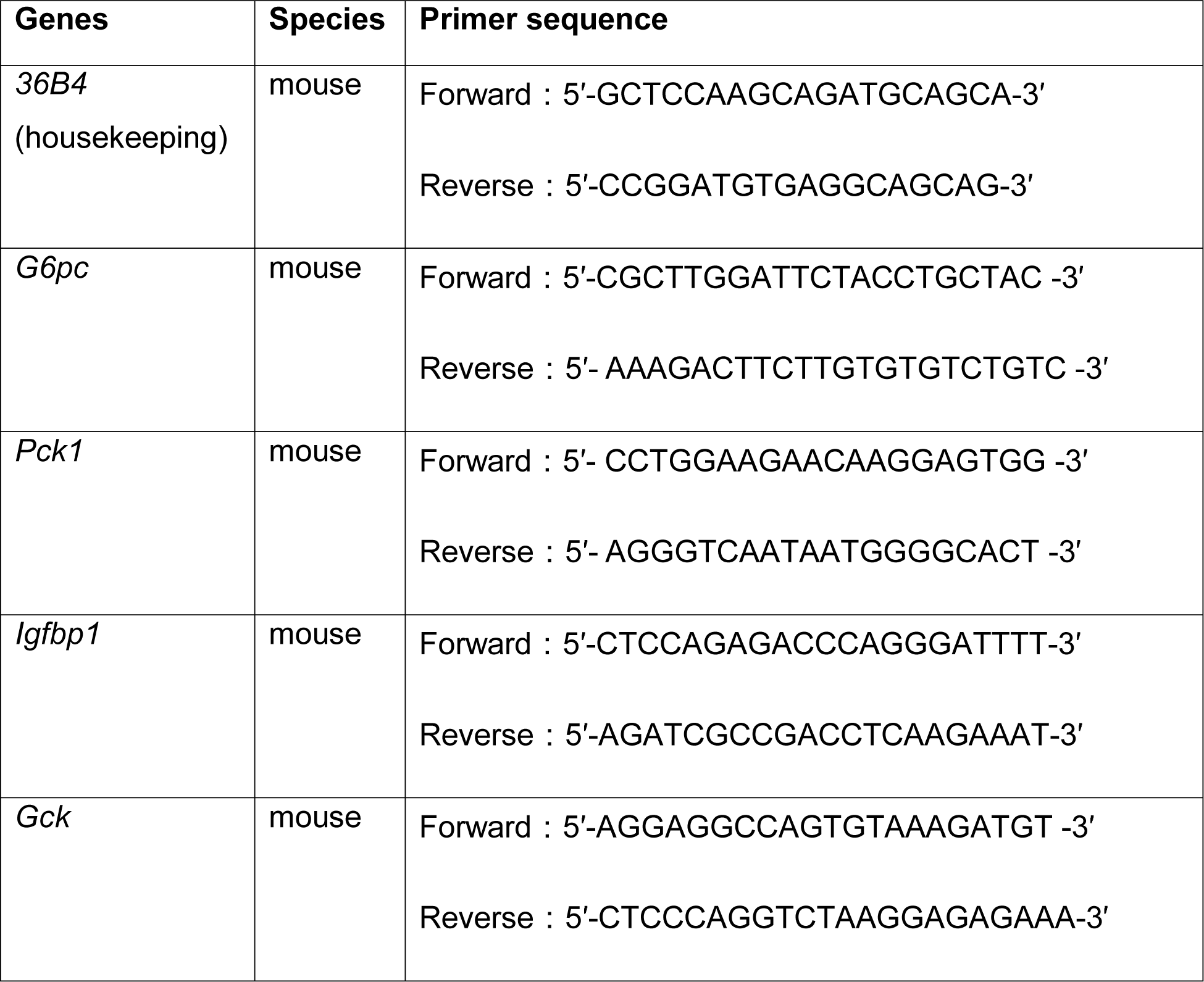
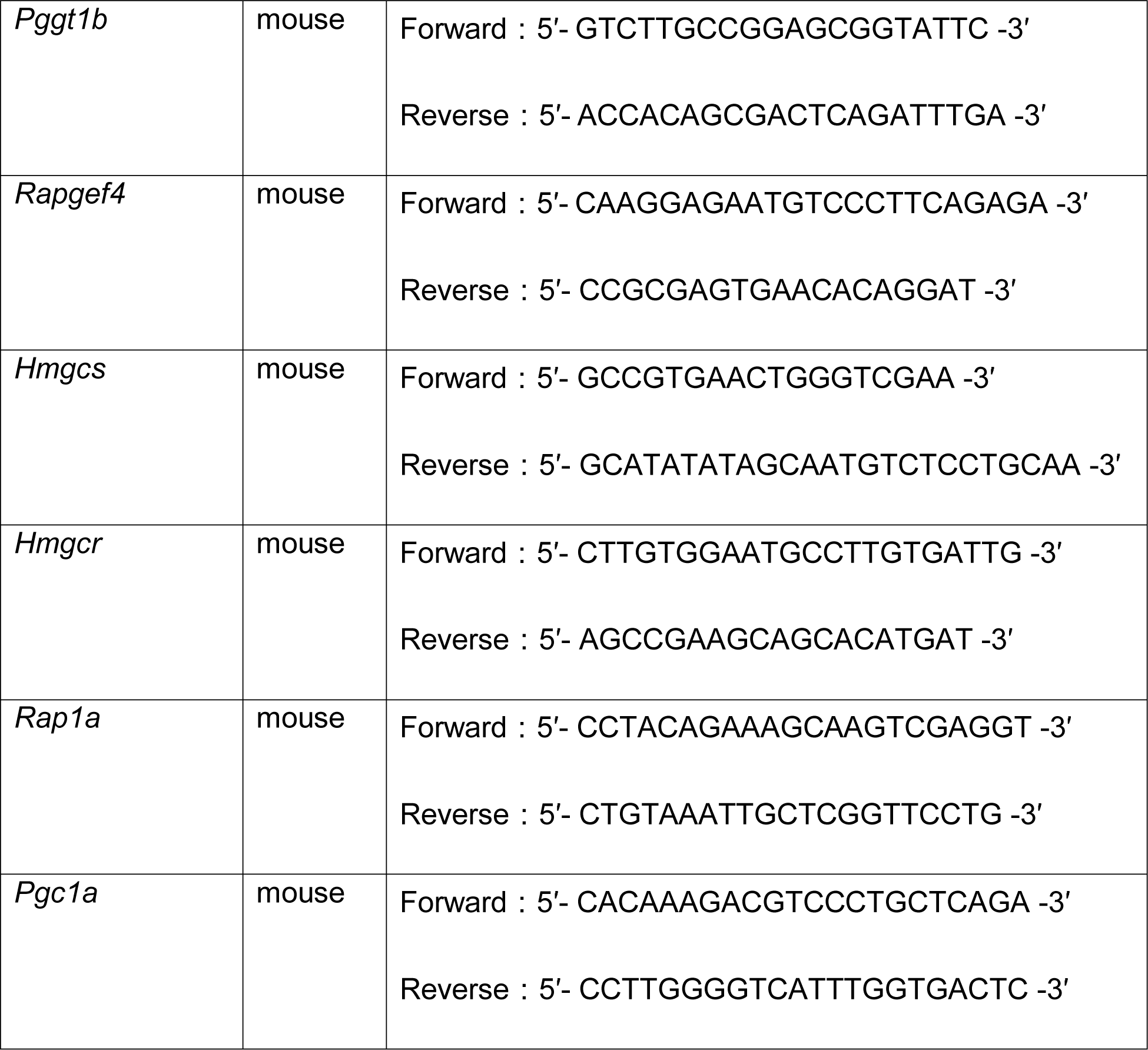
the primer sequences are designed as following sets in PCR reactions

### Statistics

All results are presented as mean ± SEM. Statistical significance was determined using SigmaPlot software. Data that passed the normality tests were analyzed using Student’s t-test and ANOVA. Data that were not normally distributed were analyzed using the nonparametric Mann-Whitney U test. Differences were considered statistically significant at p < 0.05.

### Resource Availability

Lead contact

Further information and requests for resources and reagents should be directed to and will be fulfilled by the lead contact, Dr. Lale Ozcan.

### Materials availability

This study did not generate new unique reagents.

### Data and code availability

All data reported in this paper will be shared by the lead contact upon request. This paper does not report original code.

Any additional information required to reanalyze the data reported in this paper is available from the lead contact upon request

## SUPPLEMENTAL INFORMATION TITLES and LEGENDS

**Figure S1.**
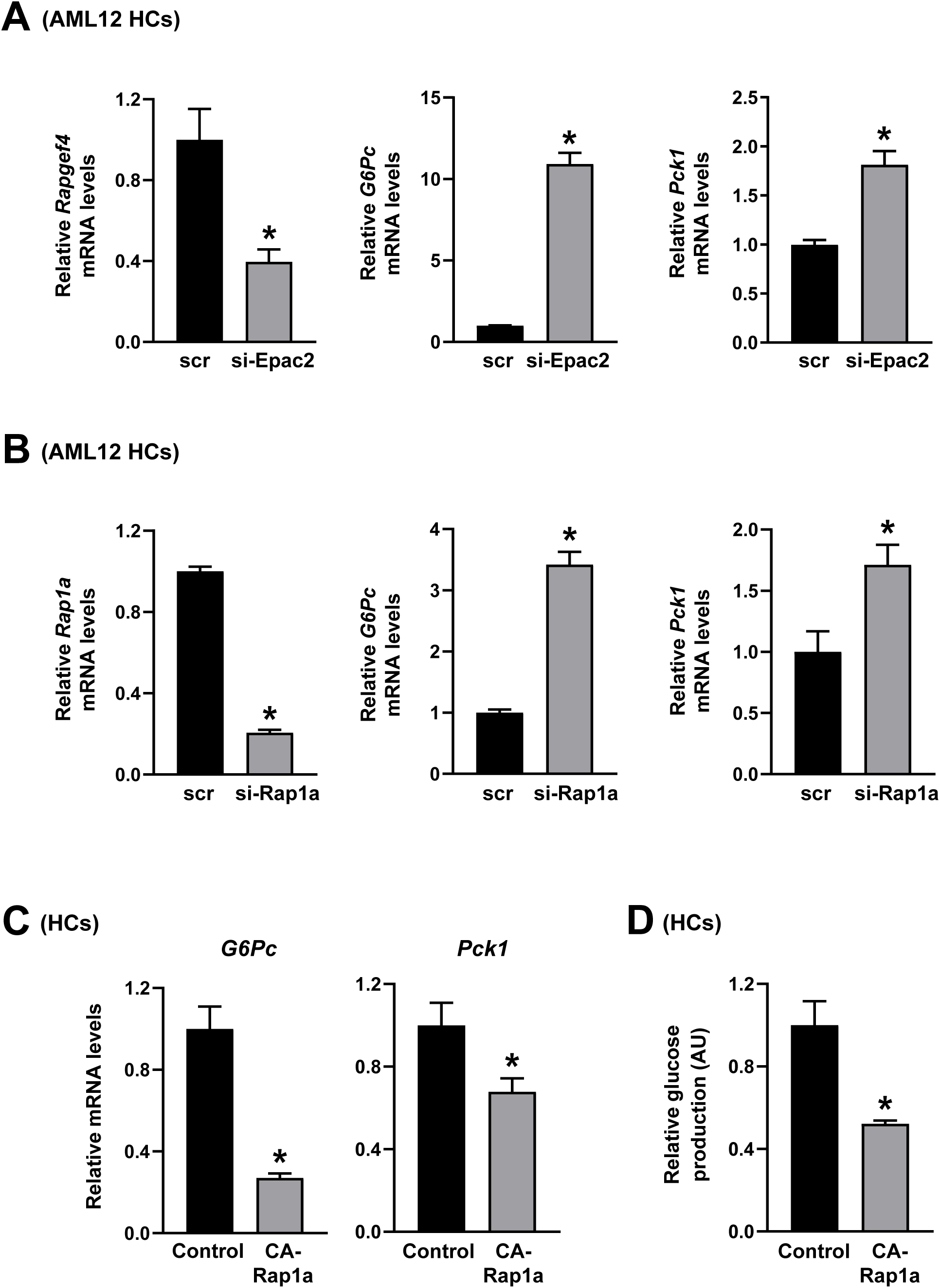
Gluconeogenesis is Induced by Epac2 or Rap1a Silencing and Suppressed by CA-Rap1a Overexpression; Related to Figure 1. **(A)** *Rapgef4* (Epac2), *G6Pc* and *Pck1* mRNA levels were analyzed from forskolin and dexamethasone-treated AML12 hepatocytes (AML12 HCs) that were transfected with scrambled RNA (scr) or siRNA against Epac2 (si-Epac2) (n= 4 wells of cells/group, mean ± SEM, *p < 0.05). **(B)** *Rap1a*, *G6Pc* and *Pck1* mRNA levels were analyzed from forskolin and dexamethasone-treated AML12 hepatocytes that were transfected with scrambled RNA (scr) or siRNA against Rap1a (si-Rap1a) (n= 4 wells of cells/group, mean ± SEM, *p < 0.05). **(C-D)** Primary mouse hepatocytes transfected with a plasmid encoding CA-Rap1a or control plasmid were treated with forskolin and dexamethasone. *G6Pc* and *Pck1* mRNA levels were analyzed (C) and glucose production was measured (D) (n= 3-5 wells of cells/group, mean ± SEM, *p < 0.05).

**Figure S2.**
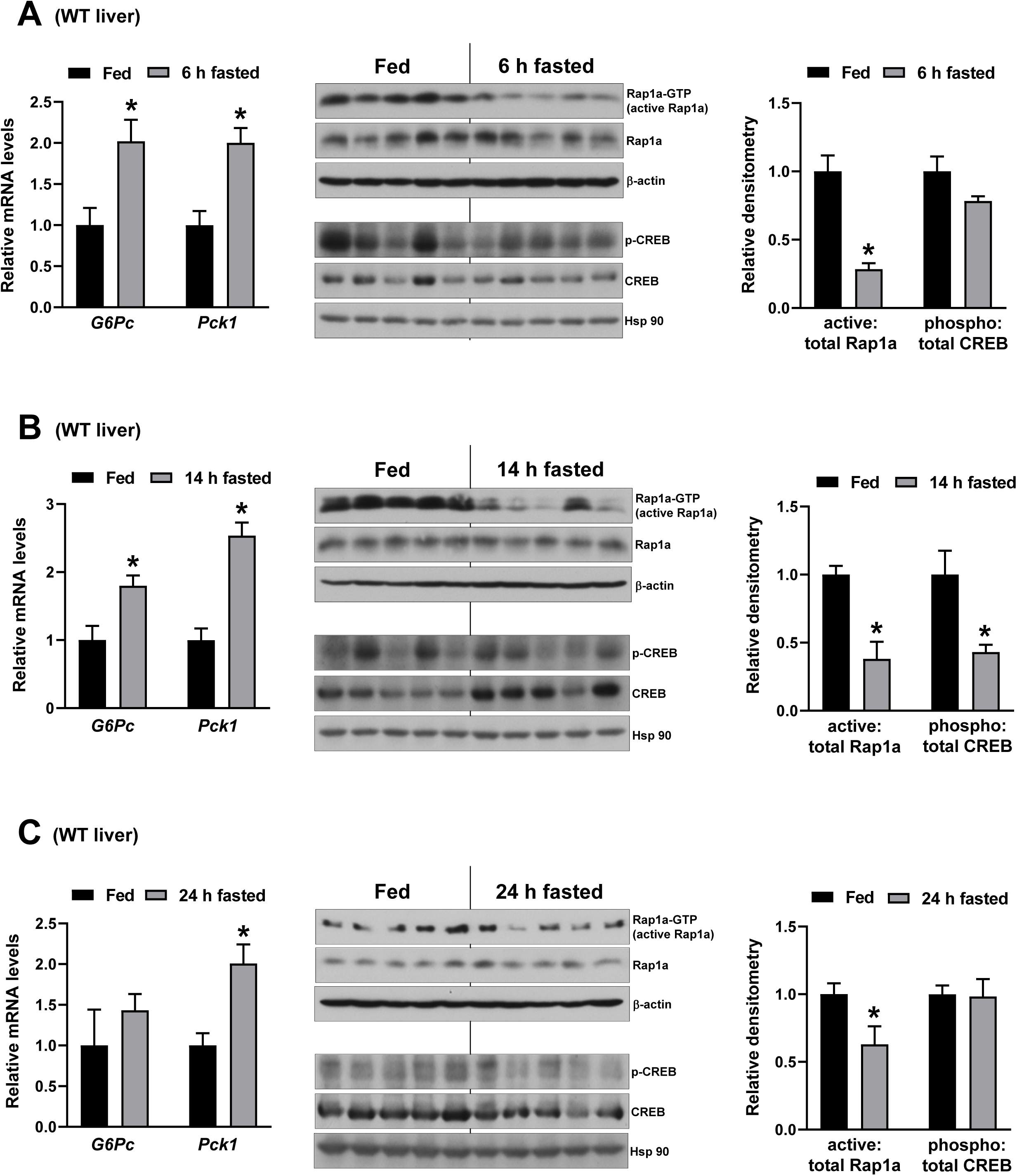
Fasting Lowers Rap1a Activity and Increases Gluconeogenic Gene Expression in WT Mice Liver; Related to Figure 1. **(A)** 16-week-old WT mice were fed ad libitum or fasted for 6 h. Hepatic *G6Pc* and *Pck1* mRNA levels were measured and active-Rap1a, Rap1a, β-actin, p-CREB, CREB and Hsp90 levels were assayed. Densitometric quantification of the immunoblot data are shown in the graph (n= 5 mice/group, mean ± SEM, *p < 0.05). **(B-C)** Same as in (A) except that mice were fed ad libitum or fasted for 14 h (B) or 24 h (C) graph (n= 5 mice/group, mean ± SEM, *p < 0.05).

**Figure S3.**
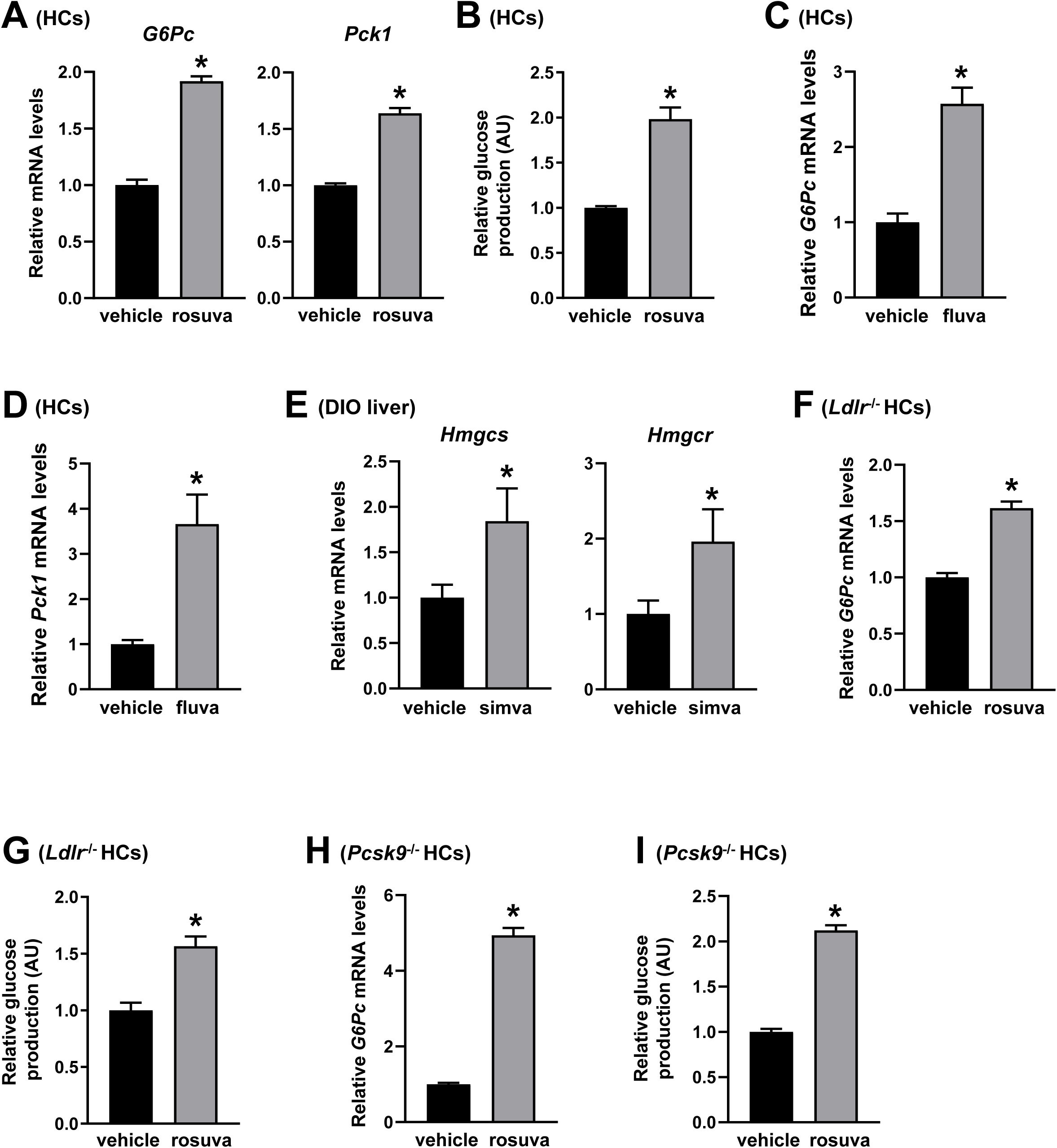
Statin Treatment Results in Increased Gluconeogenic Gene Expression in WT, *Ldlr*^-/-^ or *Pcsk9*^-/-^ Hepatocytes; Related to Figure 4. **(A-B)** Primary mouse hepatocytes (HCs) were treated with vehicle or 5 μM rosuvastatin (rosuva) for 24 h and assayed for forskolin and dexamethasone-induced *G6Pc* and *Pck1* mRNA (A) and glucose production (B) (n= 3 wells of cells/group, mean ± SEM, *p < 0.05). **(C-D)** Primary mouse hepatocytes were treated with vehicle or 10 μM fluvastatin (fluva) for 24 h and assayed for forskolin and dexamethasone-induced *G6Pc* (C) and *Pck1* (D) mRNA (n= 3 wells of cells/group, mean ± SEM, *p < 0.05). **(E)** Hepatic *Hmgcs* and *Hmcgr* mRNA levels from DIO mice that were fed with a high-fat diet containing 0.02 % simvastatin (w/w) or control high-fat diet for 12 weeks (n= 7 mice/group, mean ± SEM, *p < 0.05). **(F-G)** *Ldlr*^-/-^ primary hepatocytes were treated with vehicle or 5 μM rosuvastatin (rosuva) for 24 h and assayed for forskolin and dexamethasone-induced *G6Pc* (F) and glucose production (G) (n= 3-4 wells of cells/group, mean ± SEM, *p < 0.05). **(H-I)** Same as in (F-G) except that *Pcsk9*^-/-^ primary hepatocytes were used (n= 3-4 wells of cells/group, mean ± SEM, *p < 0.05).

**Figure S4.**
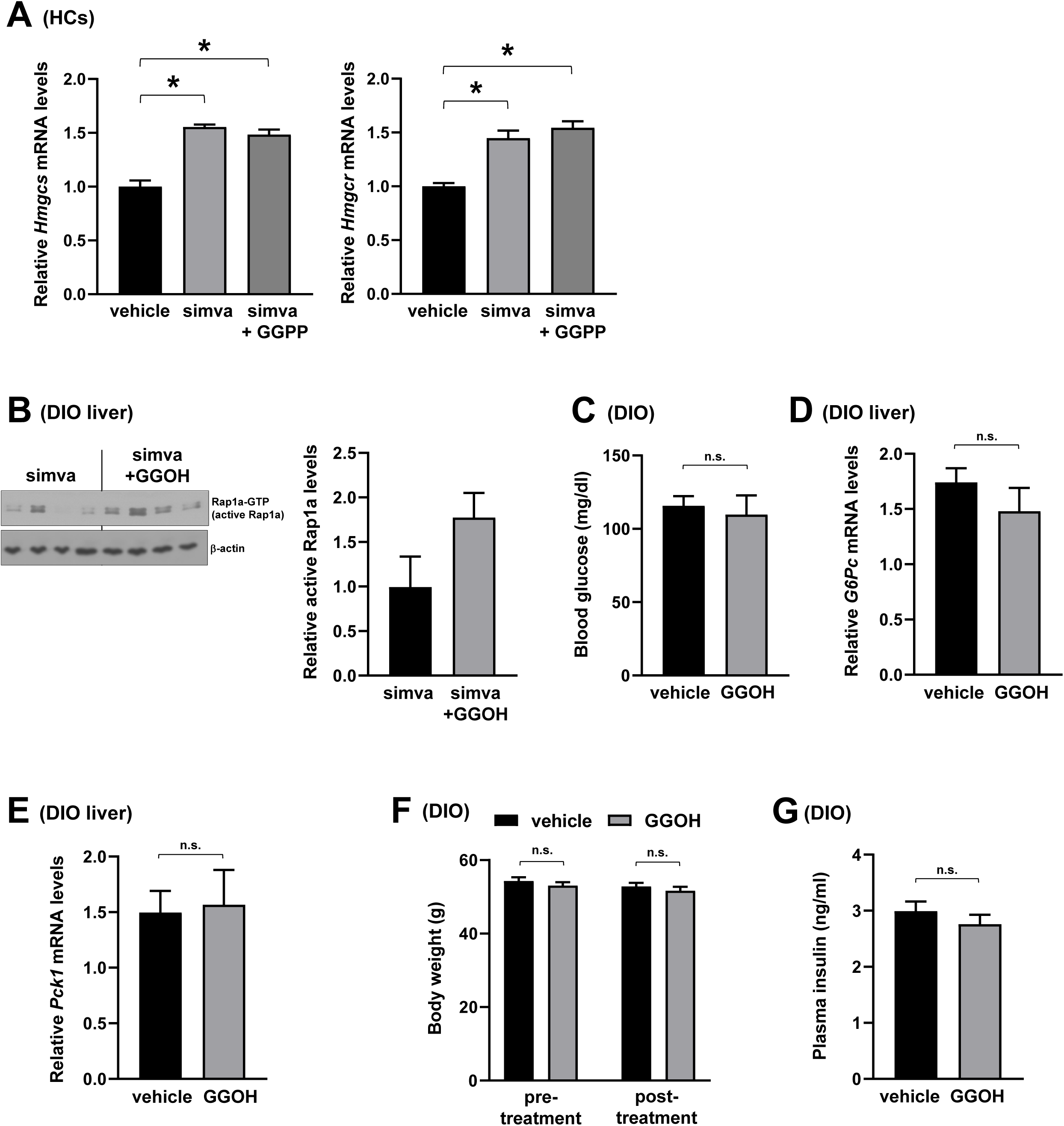
Effect of GGOH Treatment on Statin-Induced Hepatic *Hmgcs* and *Hmgcr* mRNA Levels, Rap1a Activity and Metabolic Parameters in DIO mice; Related to Figure 5. **(A)** *Hmgcs* and *Hmgcr* mRNA levels were analyzed from primary mouse hepatocytes that were treated with vehicle, 10 μM simvastatin (simva) or simvastatin + geranylgeranyl pyrophosphate (GGPP) (10 μM) for 24 h (n= 3 wells of cells/group, mean ± SEM, *p < 0.05). **(B)** DIO mice were fed with 0.02 % simvastatin (w/w)-containing high-fat diet for 12 weeks. Mice were then administered with geranylgeraniol (GGOH, 100 mg/kg/day), or vehicle control by daily gavage for 3 weeks while still receiving the statin-containing diet. Livers were assayed for GTP-bound (active) Rap1a and β-actin. Densitometric quantification of the immunoblot data is shown in the bar graph. **(C-G)** DIO mice were administered with geranylgeraniol (GGOH, 100 mg/kg/day) or vehicle control by daily gavage for 3 weeks. Overnight fasting blood glucose (C), liver *G6Pc* (D) and *Pck1* mRNA expression levels (E), body weight before and after GGOH treatment (F), and fasting insulin levels (G) were measured (n= 6 mice/group, mean ± SEM, *p < 0.05, n.s., non-significant).

**Figure S5.**
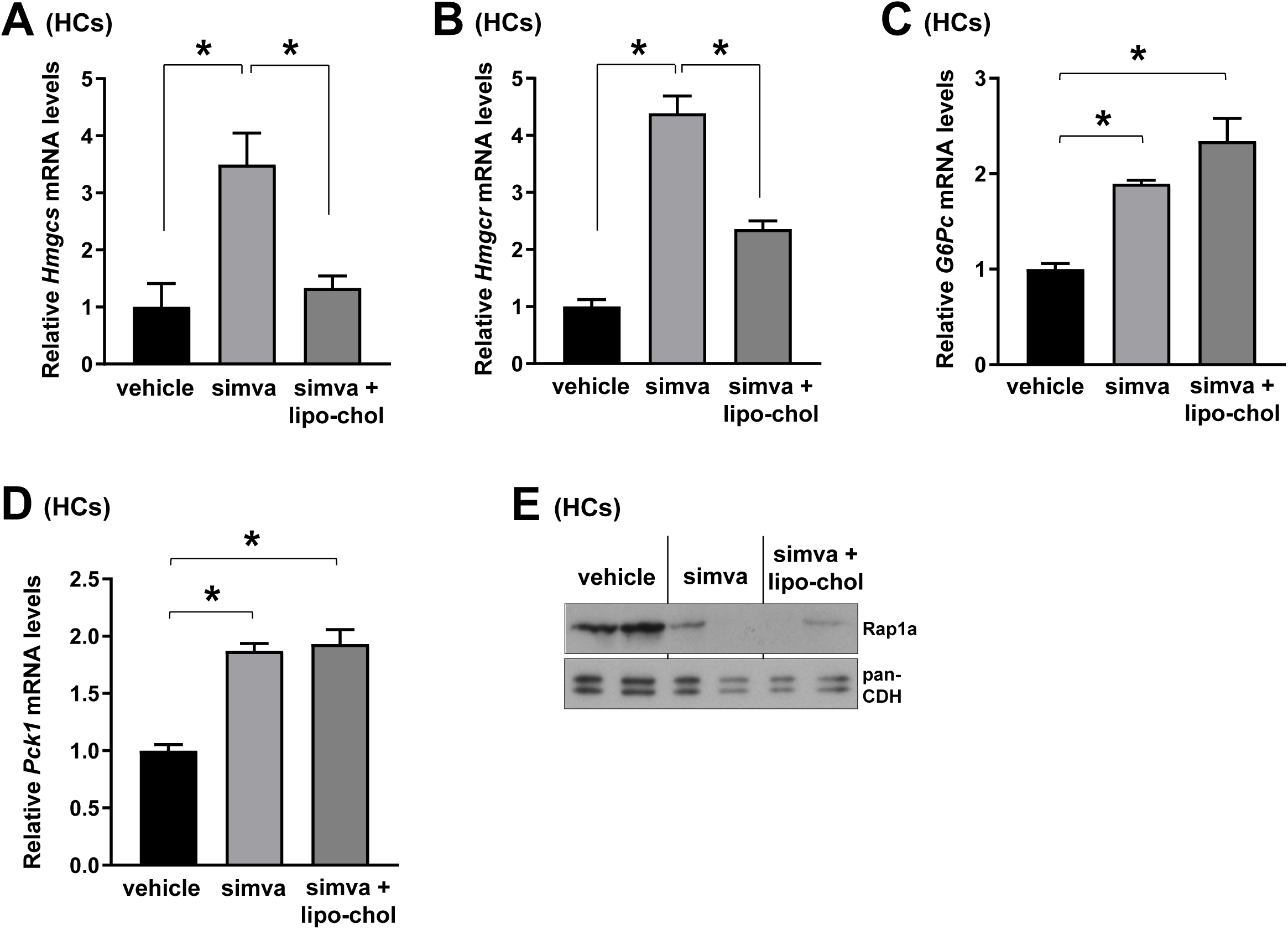
Restoration of Intracellular Cholesterol Does Not Lower Statin-Induced Gluconeogenesis; Related to Figure 5. **(A-E)** Primary mouse hepatocytes (HCs) were treated with vehicle, simvastatin (simva) or simvastatin + cholesterol-rich liposomes (lipo-chol) for 24 h and assayed for *Hmgcs* (A) and *Hmgcr* mRNA (B), forskolin and dexamethasone-induced *G6Pc* (C) and *Pck1* mRNA (D), and plasma membrane Rap1a and pan-Cadherin (pan-CDH, loading control) (E) (n= 2-4 wells of cells/group, mean ± SEM, *p < 0.05).

**Figure S6.**
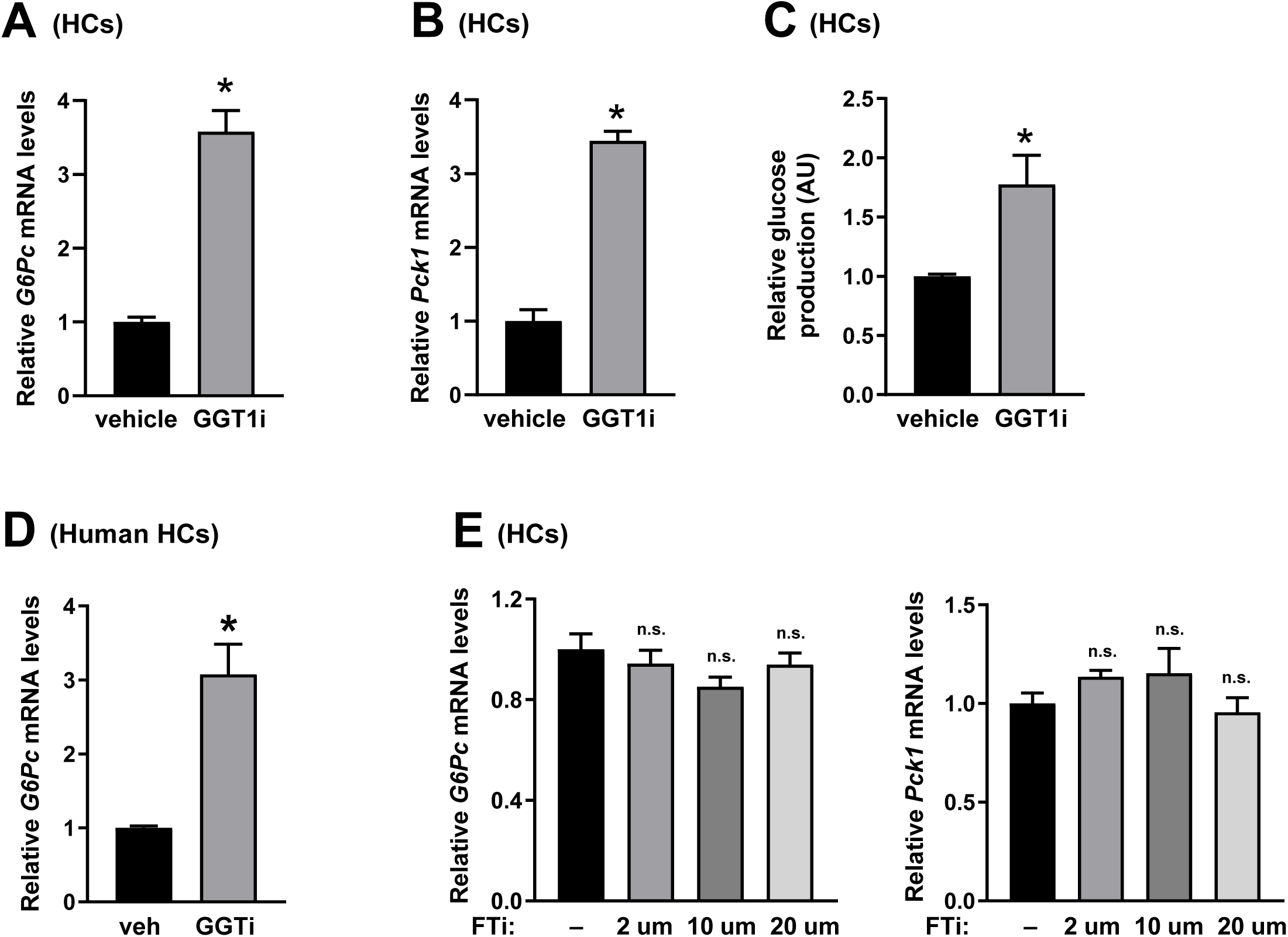
Inhibition of Geranylgeranyl Transferase, But Not Farnesyl Transferase, Results in Increased Gluconeogenic Gene Expression; Related to Figure 5. **(A-C)** Primary mouse hepatocytes (HCs) were treated with vehicle (control) or 10 μM GGT1 inhibitor (GGT1-i) for 24 h and assayed for forskolin and dexamethasone-induced *G6Pc* (A) and *Pck1* mRNA (B) and glucose production (C) (n= 3 wells of cells/group, mean ± SEM, *p < 0.05). **(D)** Primary human hepatocytes were treated with vehicle control or 10 μM GGT1 inhibitor for 24 h (GGT1i) and assayed for forskolin and dexamethasone induced *G6Pc* mRNA levels (n= 4 wells of cells/group, mean ± SEM, *p < 0.05). **(E)** Primary mouse hepatocytes were treated with vehicle control or indicated concentrations of farnesyl transferase inhibitor (FTi) for 24 h and assayed for forskolin and dexamethasone induced *G6Pc* and *Pck1* mRNA levels (n= 3 wells of cells/group, mean ± SEM, n.s., non-significant).

**Figure S7.**
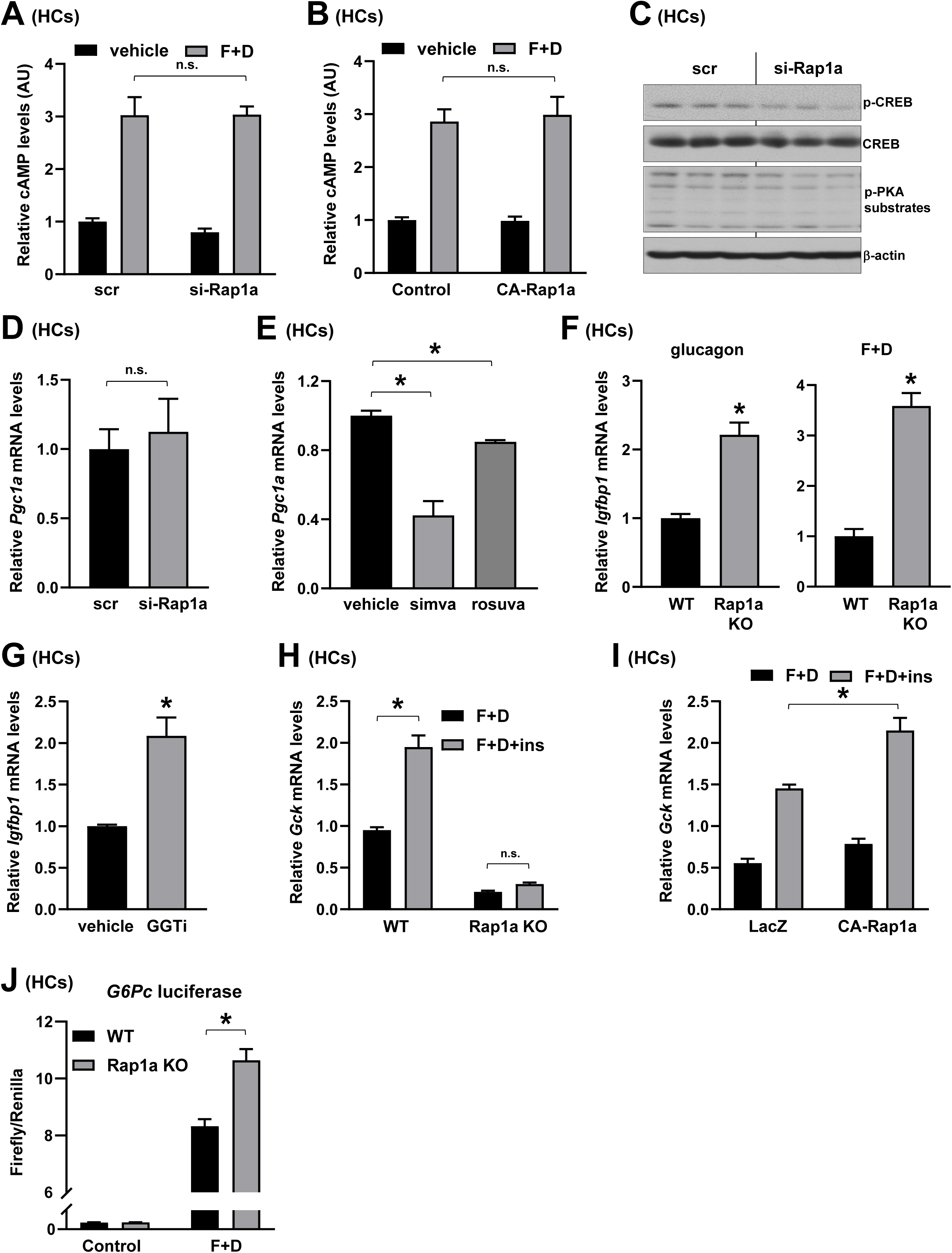
Rap1a Inhibition or Statin Treatment Regulate FoxO1 Activity Without Altering Intracellular cAMP or PKA Activity; Related to Figure 6. **(A-B)** Vehicle- and forskolin and dexamethasone (F+D)-induced intracellular cAMP levels were measured from primary hepatocytes (HCs) that were transfected with scrambled RNA (scr) or siRNA against Rap1a (si-Rap1a) (A), or a plasmid encoding CA-Rap1a or control plasmid (B) (n= 3 wells of cells/group, mean ± SEM, n.s., non-significant). **(C)** Forskolin and dexamethasone-induced p-CREB, CREB, p-PKA substrates and β-actin levels were assayed from primary hepatocytes that were transfected with scrambled RNA (scr) or siRNA against Rap1a (si-Rap1a) (n= 3 wells of cells/group, mean ± SEM, *p < 0.05). **(D-E)** Forskolin and dexamethasone-induced *Pgc1a* mRNA levels were measured from primary hepatocytes transfected with scrambled RNA (scr) or siRNA against Rap1a (si-Rap1a) (D), or primary hepatocytes treated with 10 μM simvastatin (simva) or 5 μM rosuvastain (rosuva) for 24 h (E) (n= 5 wells of cells/group and n= 3 wells of cells/group, respectively, mean ± SEM, *p < 0.05, n.s., non-significant). **(F)** *Igfbp1* mRNA levels were analyzed from WT and Rap1a KO primary hepatocytes incubated with glucagon (left panel) or forskolin and dexamethasone (F+D) (right panel) (n= 6 wells of cells/group, n= 6 wells of cells/group, respectively, mean ± SEM, *p < 0.05). **(G)** *Igfbp1* mRNA levels were analyzed from 10 μM GGTi-treated WT primary hepatocytes that were stimulated with forskolin and dexamethasone (n= 3 wells of cells/group, mean ± SEM, *p < 0.05). **(H)** *Gck* mRNA levels were measured from WT and Rap1a knock-out (KO) primary hepatocytes treated with forskolin plus dexamethasone (F+D) or forskolin plus dexamethasone plus insulin (F+D+ins) for 6 h (n= 6 wells of cells/group, mean ± SEM, *p < 0.05, n.s., non-significant). **(I)** Same as in (G) except that WT mouse hepatocytes transfected with plasmids encoding CA-Rap1a or control plasmid were used (n= 4 wells of cells/group, mean ± SEM, *p < 0.05). **(J)** WT and Rap1a KO primary hepatocytes were transfected with a luciferase fusion construct encoding nucleotides −1227 to +57 of the *G6pc* promoter containing an intact FoxO binding site. Relative luciferase activity was measured following treatment with vehicle (control) or forskolin and dexamethasone (F+D) (n= 3 wells of cells/group, mean ± SEM, *p < 0.05).

